# Evolution of genome fragility enables microbial division of labor

**DOI:** 10.1101/2021.06.04.447040

**Authors:** E.S. Colizzi, B. van Dijk, R.M.H. Merks, D.E. Rozen, R.M.A. Vroomans

**Affiliations:** Sainsbury Laboratory, Cambridge University, United Kingdom; Department of Microbial Population Biology, Max Planck Institute for Evolutionary Biology, Plön, Germany; Mathematical Institute, Leiden University; Institute of Biology, Leiden University; Origins Center, the Netherlands; Institute of Biology, Leiden University, the Netherlands

## Abstract

Division of labor can evolve when social groups benefit from the functional specialization of its members. Recently, a novel means of coordinating division of labor was found in the antibiotic-producing bacterium *Streptomyces coelicolor*, where functionally specialized cells are generated through large-scale genomic re-organization. Here, we investigate how the evolution of a genome architecture enables such mutation-driven division of labor, using a multi-scale mathematical model of bacterial evolution. We let bacteria compete on the basis of their antibiotic production and growth rate in a spatially structured environment. Bacterial behavior is determined by the structure and composition of their genome, which encodes antibiotics, growth-promoting genes and fragile genomic loci that can induce chromosomal deletions. We find that a genomic organization evolves that partitions growth-promoting genes and antibiotic-coding genes to distinct parts of the genome, separated by fragile genomic loci. Mutations caused by these fragile sites mostly delete growth-promoting genes, generating antibiotic-producing mutants from non-producing (and weakly-producing) progenitors, in agreement with experimental observations. Mutants protect their colony from competitors but are themselves unable to replicate. We further show that this division of labor enhances the local competition between colonies by promoting antibiotic diversity. These results show that genomic organization can co-evolve with genomic instabilities to enable reproductive division of labor.

**Motivation of current work:** Division of labor can evolve if trade-offs are present between different traits. To organize a division of labor, the genome architecture must evolve to enable differentiated cellular phenotypes. Cell differentiation may be coordinated through gene regulation, as occurs during embryonic development. Alternatively, when mutation rates are high, mutations themselves can guide cell and functional differentiation; however, how this evolves and is organized at the genome level remains unclear. Here, using a model of antibiotic-producing bacteria based on multicellular Streptomyces, we show that if antibiotic production trades off with replication, genome architecture can evolve to support a mutation-driven division of labor. These results are consistent with recent experimental observations and may underlie division of labor in many bacterial groups.

## Introduction

Multicellular organisms provide a clear example of a reproductive division of labor, where the germline produces gametes that generate offspring, while the disposable somatic tissues carry out functions that improve survival. Similar divisions of labor are found in colonies of social insects, where one or few individuals are responsible for all of the reproduction, whereas the other individuals perform tasks that mirror those of somatic cells in multicellular organisms. Recently, several striking examples of reproductive and other divisions of labor have been described in the microbial world [1–5]; it has been proposed that reproductive division of labor existed already before the Origin of Life, among prebiotic replicators [6–8]. Thus, such divisions of labor are ubiquitous, and the mechanisms driving them may be diverse.

In a multicellular reproductive division of labor, somatic cells typically carry the same genetic information as the germline. Cell specialization is brought about by a combination of gene regulation and epigenetics, ensuring that only a small subset of the genome is expressed. Recently, an alternative route to reproductive division of labor has been observed in the antibiotic-producing bacterium *Streptomyces coelicolor* - which generates somatic cells through mutations rather than gene regulation. Here, we propose that the genome of *S. coelicolor* has become structured over evolutionary time such that mutations occur frequently to yield differentiated cells, giving rise to a reproducible division of labor.

Streptomycetes are multicellular bacteria that grow from haploid spores, first producing a vegetative mycelium, and then differentiating into aerial hyphae that produce environmentally resistant spores. During differentiation into aerial hyphae, colonies produce a diverse repertoire of secondary metabolites [9], including antibiotics that are used to regulate competitive interactions between strains [10]. Recent results suggest that antibiotic production and spore formation are carried out by distinct cell types. Antibiotic synthesis and secretion are metabolically expensive tasks that trade off with replication [11]; accordingly, colony fitness is expected to be higher when these tasks are partitioned into separate cells [12]. The antibiotic-hyperproducing subset of cells in *S. coelicolor* colonies arises due to massive and irreversible deletions at the left and right arms of the *Streptomyces* linear chromosome [11]. Cells with larger deletions produce more antibiotics, but also produce significantly fewer spores, a deficit that effectively ensures their elimination during each replicative cycle [13]. They are instead repeatedly re-generated independently in each colony following spore germination. This process gives rise to heterogeneous colonies containing a diversity of mutants with different chromosome sizes that produce different combinations of antibiotics, as well as a larger fraction of cells specialized for spore production [11].

The irreversible mutational mechanism used to generate division of labor in *S. coelicolor* may be widespread in the genus, which is well known for its genome instability [14–16]. Many bacterial genomes are organized such that some regions of the chromosome show higher genetic variation than others, for example through the targeted insertion of mobile genetic elements, that can also carry functional genes [17]. In *Streptomyces*, these regions are located towards the telomeres of their linear chromosome, and are known to evolve rapidly through DNA amplification, deletion, recombination and conjugation [18–20]. In particular, megabase long deletions occur frequently in the Streptomyces chromosome [14, 21, 22, 11, 13]. But how does this instability reliably generate antibiotic-producing cells? Here, we hypothesize that chromosomal gene order has evolved such that some functional groups of genes have localized at the telomeric ends of the chromosome, making them more susceptible to deletion due to genome instability. By this argument, genome instability becomes adaptive within the context of this genome organization, because it facilitates the generation of sterile antibiotic-producing mutants from replicating cells. We show that a genome architecture capable of generating mutation-driven division of labor evolves in a mathematical model of antibiotic-producing bacterial colonies.

## Results

### Model overview

We formulated a mathematical model based on a simplified description of the multicellular life cycle and ecology of *Streptomyces*. We focus on the vegetative growth stage, especially during the developmental transition to sporulation [2]. In this phase, colonies grow, interact and compete with one another for space and resources [16, 10, 23]. This growth-dominated phase is followed by a phase dominated by investment in secondary metabolism [24, 9, 25, 26]. Secreted antibiotics diffuse around the producing colony, protecting it from competing strains and allowing it to claim the space into which it can grow [27]. Finally, sporulation is induced when colonies experience resource limitation.

We model individual *Streptomyces*-like cells inhabiting a two-dimensional surface, on which they replicate and secrete antibiotics. The genome organization of these virtual cells evolves through multiple cycles of vegetative growth and sporulation. Each cycle of vegetative growth starts with a population of germinating spores. Bacteria replicate locally, into empty lattice sites in their direct neighborhood. Due to replication and limited dispersal, related bacteria remain close to one another and form colonies (Fig.1a). Colonies develop for a fixed number of time steps *τ*_*s*_ (in each time step, all lattice sites are updated in random order) - after which we assume that resources are depleted and colonies sporulate. We model sporulation by randomly selecting a small fraction *ξ* of the bacteria (each cell is chosen with the same per-capita probability), which will seed the next growth cycle. Spores do not disperse between cycles (i.e. there is no spore mixing). In practice, we end the growth cycle by killing at random a fraction 1 − *ξ* of all the cells, leaving the remaining cells to initiate the next growth cycle.

**Figure 1:**
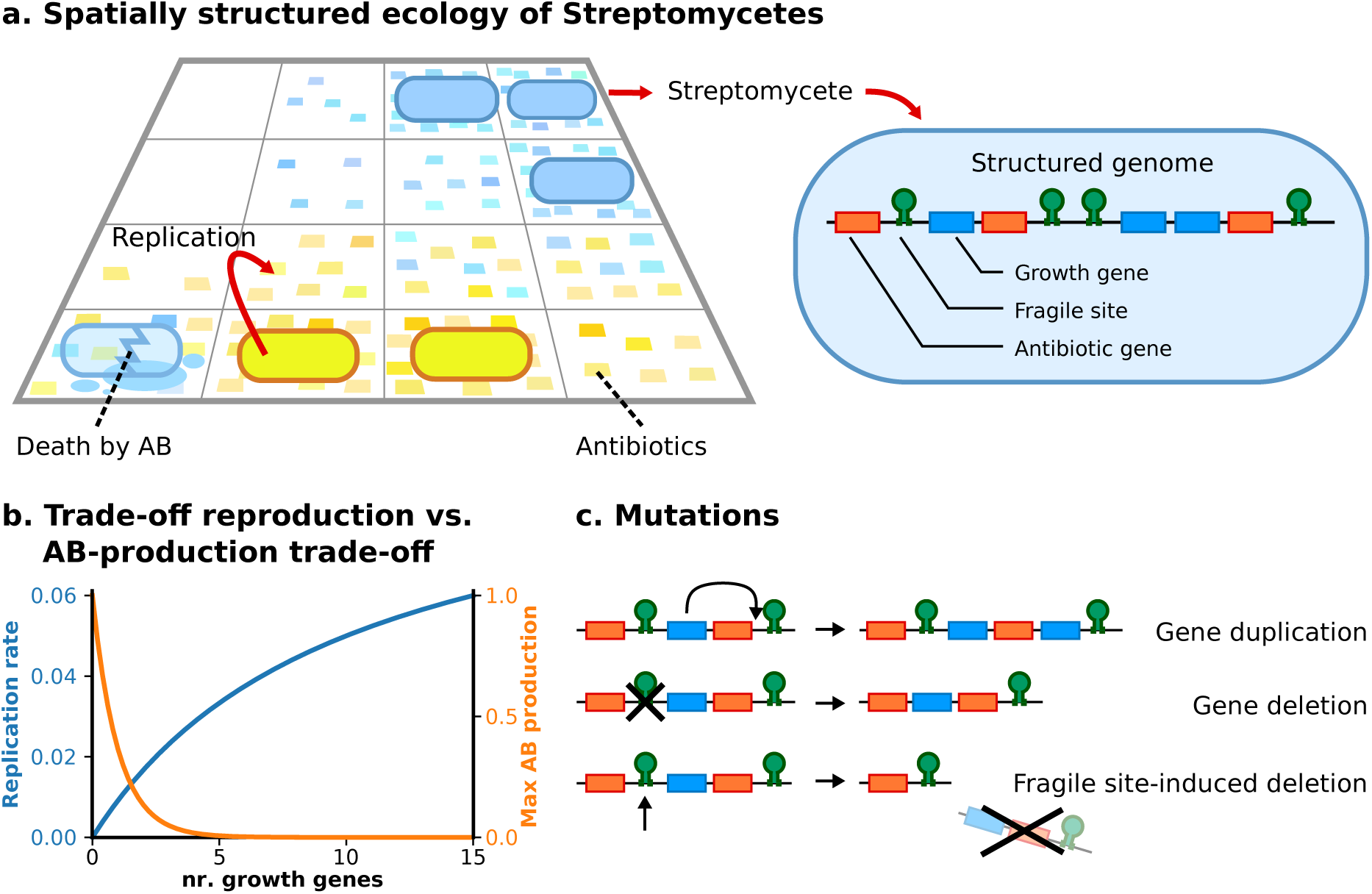
The model: a population of virtual *Streptomyces* evolving through many cycles of colonial growth of duration *τ*_*s*_ = 2500 time steps. **a** Bacteria replicate locally on a two-dimensional surface. They produce antibiotics, which are placed in their vicinity, to which other bacteria may be susceptible. Each bacterium contains a genome - a linear sequence of genes and genetic elements. We consider two gene types - growth-promoting genes and antibiotic genes - as well as fragile sites. **b** The metabolic strategy of a bacterium is determined by its genome: a larger number of growth-promoting genes translates to higher growth and lower antibiotic production. Default parameter values are: *a*_*g*_ = 0.1, *h*_*g*_ = 10, *β*_*g*_ = 1, unless explicitly stated otherwise. **c** Bacterial genomes mutate during replication: single gene duplications and deletions occur at random locations on the genome with probability *μ*_*d*_ = 10^−3^ per gene, whereas large-scale deletions occur at the genomic location of fragile-sites with probability *μ*_*f*_ = 10^−2^ per site.

Each bacterium possesses a genome that determines replication rate and antibiotic production. The model incorporates the mutational dynamics observed in the genome of *S. coelicolor*, as follows. We model the *Streptomyces* linear chromosome with a beads-on-a-string model which represents genomes as a linear sequence of genes and other genetic elements [28, 29]. In addition to growth-promoting and antibiotic-production genes, we include fragile genomic sites that are mutational hotspots. These fragile sites can represent, e.g., long inverted repeats or transposable elements, that are common within bacterial genomes [30, 31]. A genome can contain a variable number of genes and fragile sites. For simplicity, we ignore genes not directly involved in the division of labor (e.g. genes for central metabolism, homeostasis and growth-neutral genes). Because of this, a bacterium with no growth-promoting genes remains alive but it is assigned a growth rate equal to zero.

We make the simplifying assumption that the metabolic strategy of each cell, i.e., the amount of resources dedicated to growth vs. antibiotic production, is determined solely by its genotype - and ignore that these strategies may be regulated by density-dependent or other secreted cues from other bacteria [32–34]. Based on the finding that the deletion of part of the genome leads both to reduced growth and to increased antibiotic production [11], we assume that growth-promoting genes are (partly) responsible for regulating the switch from primary to secondary metabolism. In order to focus on genome architecture and its mutational consequence, we also do not explicitly include gene regulation, and assume that the metabolic switch between primary and secondary metabolism is genetically determined. Growth-promoting genes inhibit the expression of antibiotic genes (i.e. regulation is fixed), and this inhibition is lifted when growth genes are in low numbers. The model abstracts away the details of how this regulation takes place, and instead specifies a direct correspondence between the number of growth-promoting genes *g* and replication rate *k*_repl_ = *Rα*_*g*_*g/*(*g* + *h*_*g*_), and an inverse relationship between growth rate and antibiotic production *k*_ab_ = *A* exp(−*β*_*g*_*g*) (plotted in Fig.1b; *R* is a cell’s resistance to the antibiotics it is in contact with, *α*_*g*_ is the maximum replication rate, *h*_*g*_ is the number of growth-promoting genes producing half maximum growth rate, *A* is the maximum antibiotic production rate, *β*_*g*_ scales the inhibition of antibiotic production with the number of growth-promoting genes). In summary, this assumption allows us to study the genetic contributions to division of labor, and neglect regulatory contributions. It results in a trade-off because bacteria cannot simultaneously maximize both growth and antibiotic production, since the number of growth-promoting genes *g* trades antibiotic production for replication. For low values of *β*_*g*_, both growth and antibiotic production can be optimized at once, corresponding to an unrealistic situation where bacteria have arbitrary amounts of available energy. Therefore *β*_*g*_ must be sufficiently large to make the model realistic.

Bacteria can produce different antibiotics. The type of an antibiotic is determined by a bit-string of length *ν*, so the total number of possible antibiotics is 2^*ν*^. Antibiotics produced by a bacterium are secreted into its neighborhood, within a circle of radius *r*_*a*_ = 10 lattice sites - as a proxy for diffusion. Multiple types of antibiotics can be present at each lattice site (to reduce computational load, we do not consider antibiotic concentration but only its presence or absence). Bacteria are resistant to antibiotics if they encode an antibiotic-production gene with the same bit-string in their genome. Resistance decreases when the difference between the two strings increases. Cells die, and are removed from the lattice, after contact with an antibiotic to which they have no resistance. Mutations occur during cell division, when the genome is replicated, and consist of single-gene duplications and deletions with probability *μ*_*d*_ per gene, and fragile site-induced mutations that cause large deletions to the chromosome with probability *μ*_*f*_ per fragile site (Fig.1c). Novel fragile sites are spontaneously generated at random genomic locations with a small probability *μ*_*n*_ per replication. Moreover, antibiotic genes can mutate and diversify the antibiotics they encode. See Methods section for the details of the model, and Table 1 for parameter values used in all simulations (unless explicitly stated otherwise).

**Table 1:** Parameters

| Parameter | explanation | Values |
| --- | --- | --- |
| $L^2$ | lattice size | $300 \times 300$ |
| $\eta$ | neighborhood size for replication | 8 (Moore neigh.) |
| $\tau_s$ | Growth cycle duration | 2500 time steps |
| $\xi$ | Fraction of spores to seed a growth cycle | 0.001 |
| *Replication* |  |  |
| $\alpha_g$ | max replication probability per unit time | 0.1 |
| $h_g$ | nr. of growth genes for 1/2 max growth rate | 10 |
| $\beta_r$ | antibiotic resistance factor | 0.3 |
| $\mu_d$ | Duplication/deletion probability per gene | 0.001 |
| $\mu_f$ | Fragile site-induced deletion probability | 0.01 (per fragile site) |
| $\mu_n$ | Probability of new fragile site formation | 0.01 per genome |
| $\mu_a$ | Probability of antibiotic type mutation | 0.005 (per ab gene) |
| *Antibiotic production* |  |  |
| $\alpha_a$ | max antib. production probability per unit time | 1 |
| $h_a$ | nr. of antib. genes for 1/2 max production rate | 3 |
| $r_a$ | max distance of antib. placement | 10 |
| $\beta_g$ | antib. production decrease due to trade-off | 1 |
| $\mu$ | length of antib. bit-string | 16 |
| *Other parameters* |  |  |
| $p_{\text{mov}}$ | prob. of migration into empty adjacent site | 0.01 |

### Eco-evolutionary dynamics of virtual *Streptomyces*

Starting from short genomes (initial size = 10 genes) without fragile sites, containing homogeneously distributed antibiotic genes and growth-promoting genes, we let bacteria evolve over at least 500 vegetative growth cycles, each of *τ*_*s*_ = 2500 time steps. Over this time, bacteria incorporate about 10 fragile sites in their genome (Fig. 2a), evolve a large and diverse set of antibiotic genes (30 to 130 genes), and a approximately 10 growth-promoting genes (see Suppl. Section S1 for additional runs). After evolution, the spatial dynamics of the model over the course of one growth cycle qualitatively reproduce those of Streptomyces colony formation (see, e.g., pictures in [11, 27]): colonies expand and produce a growing halo of antibiotics and a corresponding zone of inhibition of competing strains. Colony expansion continues until colonies encounter antibiotics to which they are susceptible - i.e. antibiotics produced by other colonies. The invasion dynamics reach a quasi-stable spatial configuration once all colonies have become enclosed by antibiotics from competing strains. At this stage bacteria born at the edge of the colony are killed by an adjacent colony’s antibiotics [35] (Fig. 2b; for a time course see Suppl. Section S2 top panes, and Suppl. Video 1).

**Figure 2:**
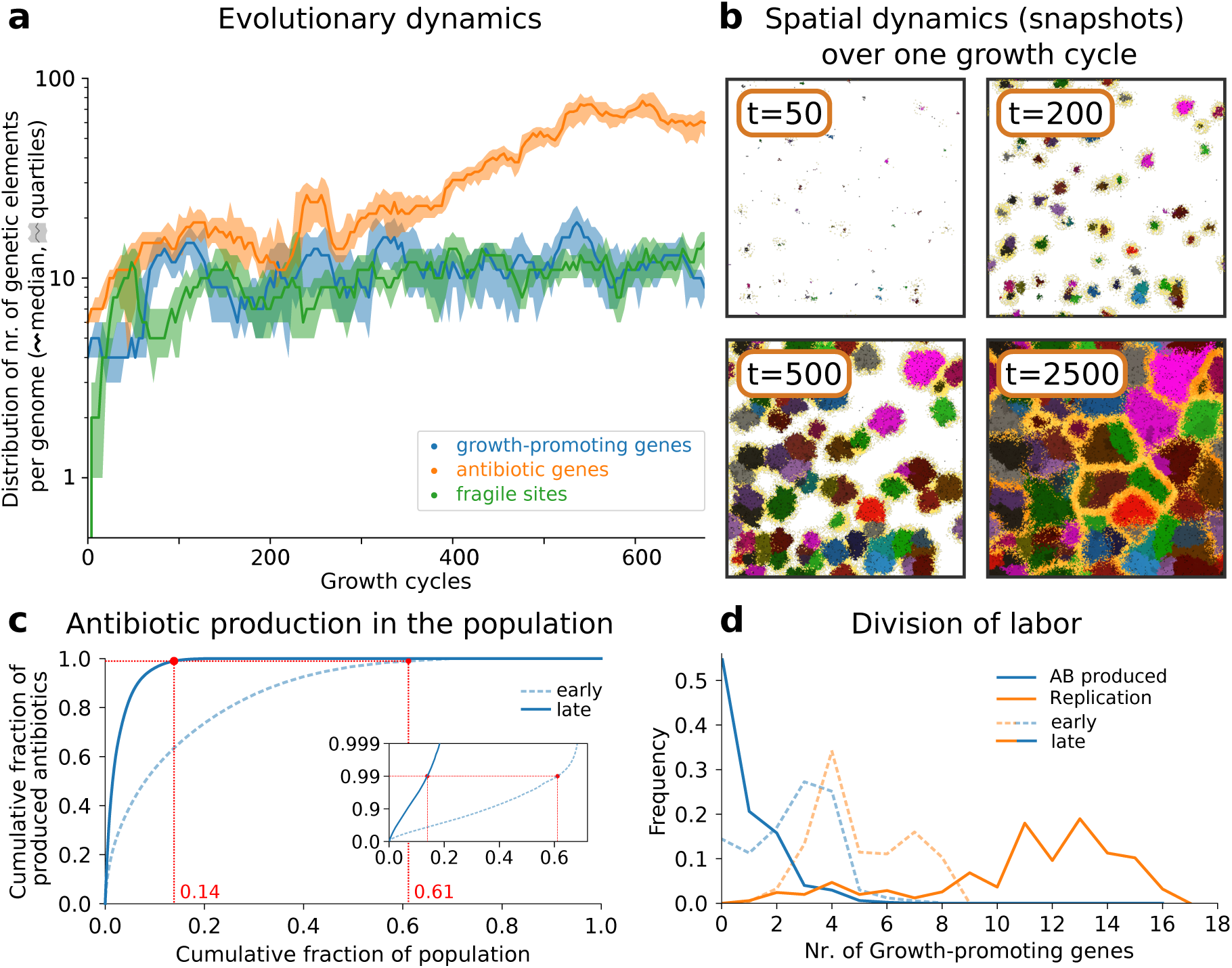
Evolution of genome composition and division of labor. **a** Evolutionary dynamics of gene content. For each time point, the median and quartile values of the number of each type of genetic element is calculated from all genomes in the population (at the end of the growth cycle). Growth cycle duration *τ*_*s*_ = 2500 time steps. Notice the logarithmic y-axis. The system is initialized with a population with genome: `5’-FAAFAAFAAF-3’`, where F is a growth gene and A is an antibiotic gene, with each antibiotic gene encoding a different antibiotic type. **b** Spatial dynamics during one growth cycle, after evolution: different colors represent different colonies, yellow shading around the colonies indicates antibiotics, time stamps in the pictures indicate the time elapsed in the growth cycle. **c** Most antibiotics are produced by few bacteria after evolution. We collect all bacteria alive at the midpoint of the growth cycle of an early (after 40 cycles, i.e. 100 *×* 10^3^ time steps) and later stage (after 400 cycles, i.e. 1000 *×* 10^3^ time steps) in the evolutionary dynamics (i.e. at 1250 time steps). For each time point, bacteria are sorted on antibiotic production, the cumulative plot (blue line) is normalized by population size and total amount of antibiotics produced, the red dot indicates the fraction of the population that produces 99% of all antibiotics. Inset: semi-log plot of the same data. **d** Division of labor between replication and antibiotic production, shown as a function of the number of growth-promoting genes. Using the same data set as **c**, the plot shows the frequency of replication events (orange) and antibiotic production events (blue) per number of growth genes, for early and later stage in evolution.

### Evolution of genome architecture leads to division of labor between replicating and antibiotic-producing bacteria

To understand the population dynamics produced by the model, we compare populations after different stages in the simulations. At an early stage (after 40 growth cycles, i.e. 100 *×* 10^3^ time steps), antibiotics are collectively produced by the whole population. By contrast, at a later stage (400 cycles, i.e., 1000 *×* 10^3^ time steps) nearly all antibiotics are produced by a small fraction of the bacteria (Fig. 2c, 99% of all antibiotics are produced by 61% of the population at time 100 *×* 10^3^, and by just 14% at time 1000 *×* 10^3^). We also observe that replication and antibiotic production are performed by genetically distinct bacteria in the later stages: bacteria with few growth-promoting genes produce most antibiotics but do not replicate frequently, whereas bacteria with a larger number of growth-promoting genes replicate frequently but do not produce antibiotics (Fig. 2d). Because antibiotic-producing bacteria do not replicate autonomously, or replicate at very low rate, they cannot form an independent lineage, and are instead generated in each growth cycle from the bacteria that have high replication rates. Therefore, antibiotic producing bacteria are found in irregular dotted pattern over the colony (Suppl. Section S2 bottom panes, and Suppl. Video 2).

We characterize the fitness cost associated to generating these mutants by performing a series of competition experiments where evolved colonies (wildtype) are pitted against generalists with a broad range of antibiotic production and growth rates. These “artificial” generalists do not pay a fitness cost for being able to replicate and produce antibiotics at the same time, and thus do not experience a trade-off. A generalist with similar growth and antibiotic production rate to a wildtype only barely outperforms the wildtype (Suppl. Section S3). This shows that the fitness cost associated with generating the mutants does not greatly affect colony competitiveness, in agreement with [11]. A steeper trade-off between replication and antibiotic production (corresponding to the metabolic shift from primary to secondary metabolism) makes the division of labor more likely to emerge. This is achieved when fewer growth-promoting genes are required to inhibit antibiotic production (i.e. lower *β*_*g*_), or when more growth-promoting genes are required for faster growth (higher *h*_*g*_; Suppl. Section S4). For *h*_*g*_*β*_*g*_ *>* 5 division of labor can evolve, and the number of genes at steady state depends on the numerical value of the two parameters (Suppl. Section S5). Higher antibiotic production rate (especially for mutants with few growth-genes) could be expected to synergize with a steeper trade-off for the evolution of division of labor. Strikingly, we find the opposite: division of labor evolves when bacteria produce fewer overall antibiotics (lower *α*_*a*_), under shallow trade-off conditions (*h*_*g*_*β*_*g*_ = 5; see Suppl. Section S6). Altogether these results suggest that the requirements for evolving division of labor in Streptomyces might be rather broad, and therefore that it is likely to evolve in species other than *S. coelicolor*: it suffices that bacteria that grow rapidly cannot meet the antibiotic production requirements for the entire colony.

### Genome architecture supports the mutation-driven division of labor

To understand the role of genome architecture in the division of labor, we extracted the founder of the most abundant colony after long-term evolution, and tracked its diversification after it was reinoculated into an empty grid (for ease of interpretation the only mutations we allow in this experiment are fragile site-induced deletions). We observed that the genome of the evolved colony founder had two distinct regions: a telomeric region that contains growth-promoting genes but lacks fragile sites, and a centromeric region that lacks growth-promoting genes and has abundant fragile sites (Fig. 3b). After colony growth from this founder, nearly all the bacteria capable of replicating are genetic copies of the founder, while most of the antibiotic production is carried out by a diverse suite of mutants arising through genomic instabilities (Fig. 3a). When bacteria divide, mutations induced at fragile sites lead to the deletion of the part of the genome distal to them, causing large telomeric deletions. Because growth-promoting genes are over-represented in these regions, they are frequently deleted as a group (Fig. 3c). Mutants generated from these deletions lack growth-promoting genes while retaining many antibiotic genes, and will therefore produce antibiotics at much higher rates (Fig. 3d). By this process, colonies that begin clonally, eventually evolve to become functionally differentiated throughout the growth cycle. On average, a colony contains 2% to 7% mutants with large telomeric deletions that specialize in antibiotic production while foregoing replication themselves (Suppl. Section S7). In Supplementary Section S8 we show this genome architecture is prevalent in the evolved population.

**Figure 3:**
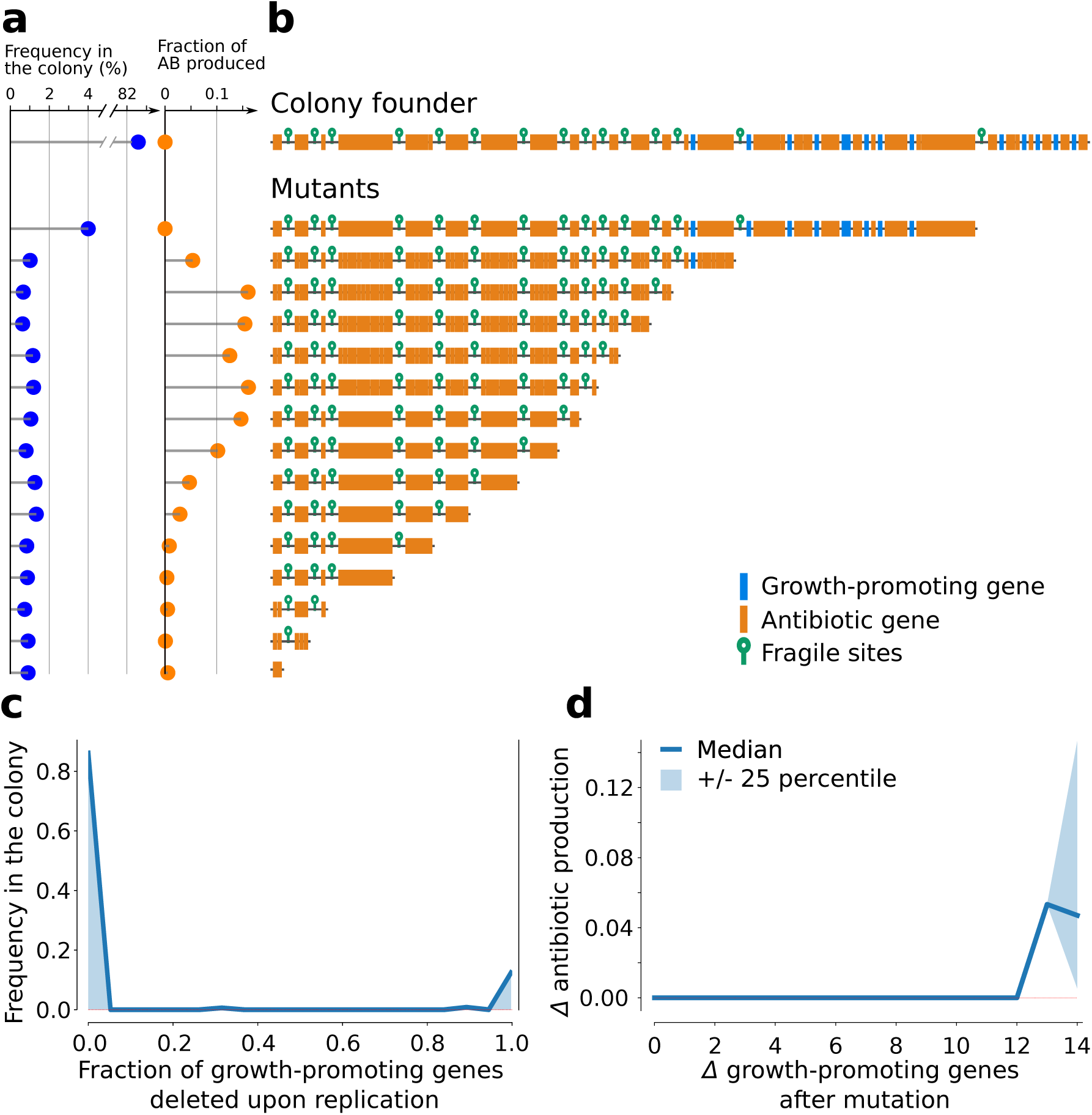
Genome architecture enables division of labor within an evolved colony: the colony founder generates antibiotic-producing somatic cells through large-scale chromosomal deletions, caused by fragile sites. The data is generated by seeding a simulation with one bacterium - the colony founder - and letting the colony grow until it reaches a diameter of 70 lattice sites (which is the approximate colony size at the end of a growth cycle, cf. Fig. 2b). **a** Genomes in high abundance do not produce many antibiotics. All bacterial genomes are collected after colony growth, they are sorted by sequence length and their centromeres are aligned to emphasize telomeric deletions. The bar-plots contrast genome frequency in the colony (blue) and the fraction of antibiotics produced by bacteria with that genome (orange). Note the broken axis for genome frequency. **b** Genome architecture of the colony founder and all its descendants. Growth-promoting genes in blue, antibiotic genes orange, fragile sites depicted as green hairpins. **c** Deletions of all-or-no growth-promoting genes during replication. Shown is the fraction of growth-promoting genes deleted because of fragile site instability upon replication during colony development (ranging from 0, i.e. no genes are deleted, and 1, all growth genes are deleted). **d** Increased antibiotic production as a result of chromosomal deletions. Difference in antibiotic production as a function of the growth-promoting genes lost during replication (colony median +*/*− 25th percentile).

To assess whether genome architecture is required for division of labor, we initiated an evolutionary run where we shuffled the order of the genes in the genome of each spore at the beginning of every growth cycle. This disrupts the genomic architecture of colony founders without changing its composition, in terms of the number and type of genes and fragile sites. Starting from an evolved genome, we observe a rapid decrease of antibiotic genes and of growth-promoting genes, resulting in small genomes (Fig.4a). At evolutionary steady state, a large fraction of the population contributes to antibiotic production (Fig.4b). Because these bacteria do not divide labor through mutation (Fig.4c), they can only express sub-optimal levels of both growth-promoting genes and antibiotic genes. In other words, they are generalists that do not resolve the trade-off, but compromise between replication and antibiotic production. Importantly, these evolved generalists always lose in direct competition experiments with bacteria that can divide labor when the genome is not mixed (Suppl. Section S9). We conclude that genome architecture is a key prerequisite for the maintenance of a mutation-driven division of labor, and therefore that genome architecture is subject to natural selection.

**Figure 4:**
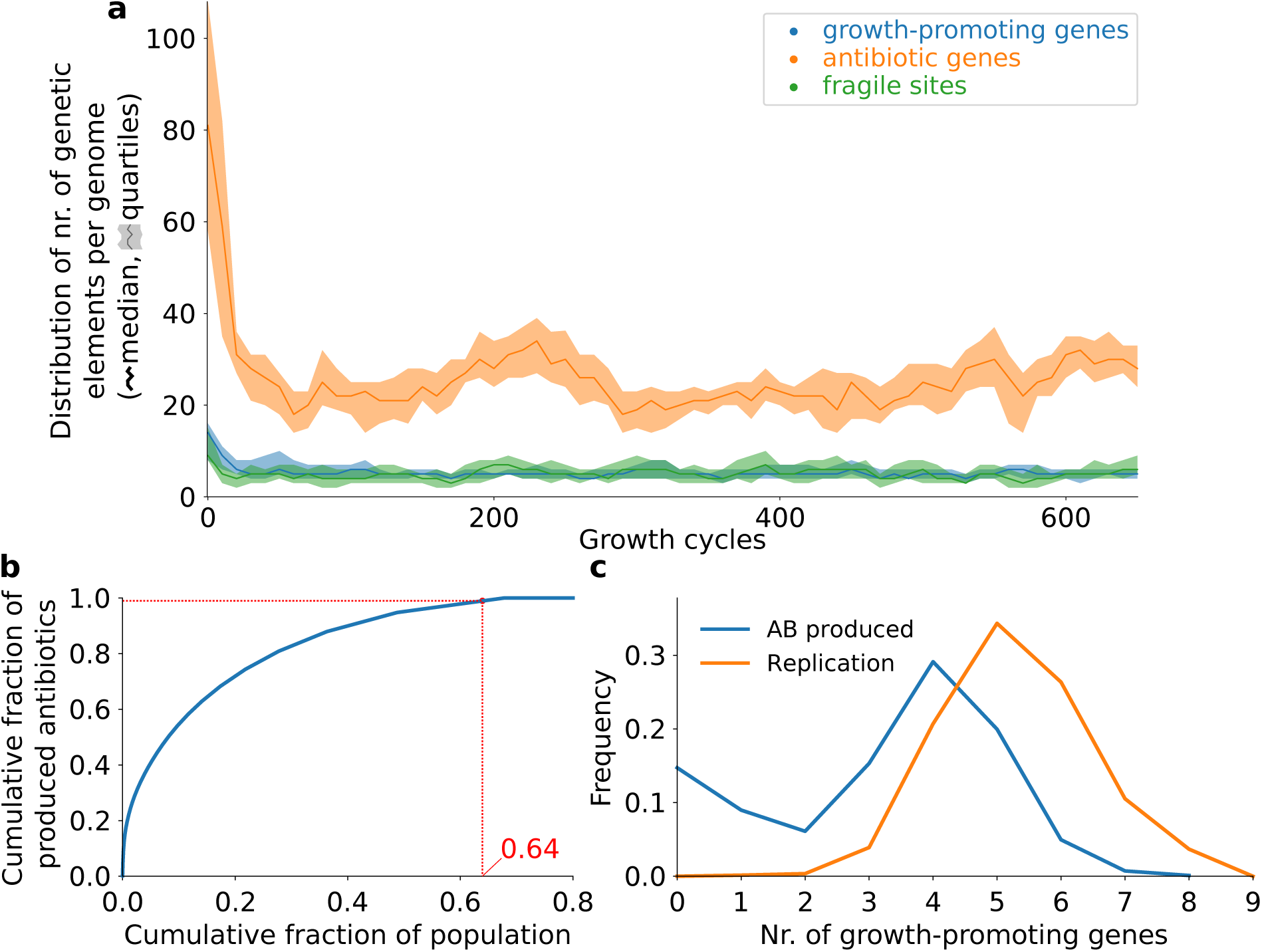
Genome shuffling between growth cycles lead to bacteria with small genomes that do not divide labor. **a** Evolutionary dynamics of the genome composition. As in Fig. 3, the median and quartile values of the number of each type of genetic element is calculated from all genomes in the population, at the end of every growth cycle. Growth cycle duration *τ*_*s*_ = 2500 time steps. We start the simulation from an evolved genome capable of mutation-driven division of labor. **b** Cumulative antibiotic production as a function of the cumulative fraction of the population. Bacteria from the mid-point of the growth cycle are sorted on antibiotic production (after evolutionary steady state is reached). The curve is normalized by population size and total amount of antibiotics produced. In red we indicate the fraction of the population that produces 99% of all antibiotics. **c** Frequency of replication (orange) and antibiotic production (blue) as a function of the number of growth-promoting genes. The same data are used as in **b**.

We also studied what genome architecture evolves when additional genes are introduced that are essential for survival but are not directly involved in the division of labor (e.g. homeostatic genes). We modeled these genes as an additional gene type, and assumed that they must be present in at least *n*_*h*_ = 10 copies for the cell to remain alive. Any number of genes lower than *n*_*h*_ is lethal and the bacterium carrying such genome is removed from the lattice, while a larger number is neutral. We find that mutation-driven division of labor evolves, and that the centromeric part of the evolved genomes contains always more than *n*_*h*_ homeostatic genes, ensuring survival of the antibiotic-producing species that arise through mutation (Suppl. Section S10).

### Effect of spatial competition dynamics and antibiotic diversity on evolution of division of labor

We next examined the effect of spatial structure on the evolution of division of labor in our model. Previous work showed that spatial structure can promote the coexistence and diversity of antibiotic producing cells, because antibiotics secreted in a cell’s neighborhood prevent invasion by adjacent strains [10, 35–39, 23]. In our model, antibiotic-based competition occurs at the boundaries of two colonies. Formation of colony boundary requires sufficiently long growth cycles (*≥* 1000 time steps; see Suppl. Section S11). Consequently, no division of labor evolves when cycles are shorter because selection only favors growth. Growth cycles shorter than 500 time steps lead to extinction as populations do not recover after sporulation. To further test the importance of competition at colony interfaces, we can perturb the spatial structure by mixing the system after each time step. Evolved bacteria maintain mutation-driven division of labor in such a mixed system (Suppl. Section S12, Fig. SF14), but division of labor cannot evolve de novo from random genomes (Fig. SF15).

We find that colonies produce a large number of different antibiotics, which results from selection for diversity (Suppl. Section S13; in line with previous work [35]). The large number of antibiotic genes in the evolved genomes is likely made possible by the simplified mutational dynamics we implemented, that allow new antibiotic types to evolve from pre-existing antibiotic genes through single mutations. When the total number of possible antibiotics is large (about 6.5 *×* 10^5^ different antibiotics, see Methods), colonies are highly susceptible to the antibiotics produced by other colonies (Suppl. Section S14). The spatial dynamics allow for the coexistence of multiple strains because they mutually repress each others’ invasion through the local production of antibiotics (Fig. SF3). With a small number of possible antibiotics, bacterial genomes evolve to contain every possible antibiotic gene, which results in low susceptibility (Suppl. Section S15, also cf. [36]). Nevertheless, mutation-driven division of labor evolves in both cases (Suppl. Section S16). Colonies also evolve division of labor when the deposition radius around the producing bacterium is much reduced (the deposition radius is a proxy for diffusion; Suppl. Section S17). This reduces the benefit afforded by antibiotics because a single antibiotic-producing bacterium can protect fewer colony members. Only for an unrealistically small antibiotic deposition radius does division of labor not evolve, and instead growth rate alone is maximized (similarly to genomes that evolve with short growth cycles, see Suppl. Section S11). The benefit of antibiotic production can also be reduced by decreasing the specificity of bacterial resistance to antibiotics (lowering *β*_*r*_, see Methods). However, division of labor evolves when specificity is decreased by more than two orders of magnitude (Suppl. Section S17).

### Fragile site mutation rates co-evolve with division of labor

Two parameters control the influx and activity of fragile sites: the rate of *de novo* fragile site formation *μ*_*n*_ and the fragile site-induced deletion probability *μ*_*f*_. We assess the robustness of our results when these two parameters are systematically varied. In fig. 5, we characterize the effect of mutation rates on the evolution of division of labor with a “division of labor index” (defined as the difference between the median number of growth-promoting genes for replicating bacteria and antibiotic-producers). A larger value indicates a larger genetic distance between the cells that replicate and produce antibiotics, and thus a stronger division of labor). We find a sharp boundary between mutation rates that allow the evolution of division of labor (blue tiles) and mutation rates for which division of labor does not evolve (red tiles; See Suppl. Section S18 for more data). The evolution of division of labor is accompanied by evolution of high rate of genome deletion (black line in Fig. 5). A larger number of fragile sites is incorporated in the genomes when each fragile site has a smaller probability of causing deletion (Suppl. Section S19). The evolution of division of labor also results in the incorporation of a larger number of growth-promoting genes. When division of labor does not evolve, genomes contain few growth-promoting genes, so that each bacterium grows and produces antibiotics. Bacteria that do not evolve division of labor behave like generalists, and are similar to those evolved with genome shuffling, see Fig. 4. However, once division of labor evolved with *μ*_*n*_ *>* 0, it persists in the evolved populations when *μ*_*n*_ is set to zero (Suppl. Section S20).

**Figure 5:**
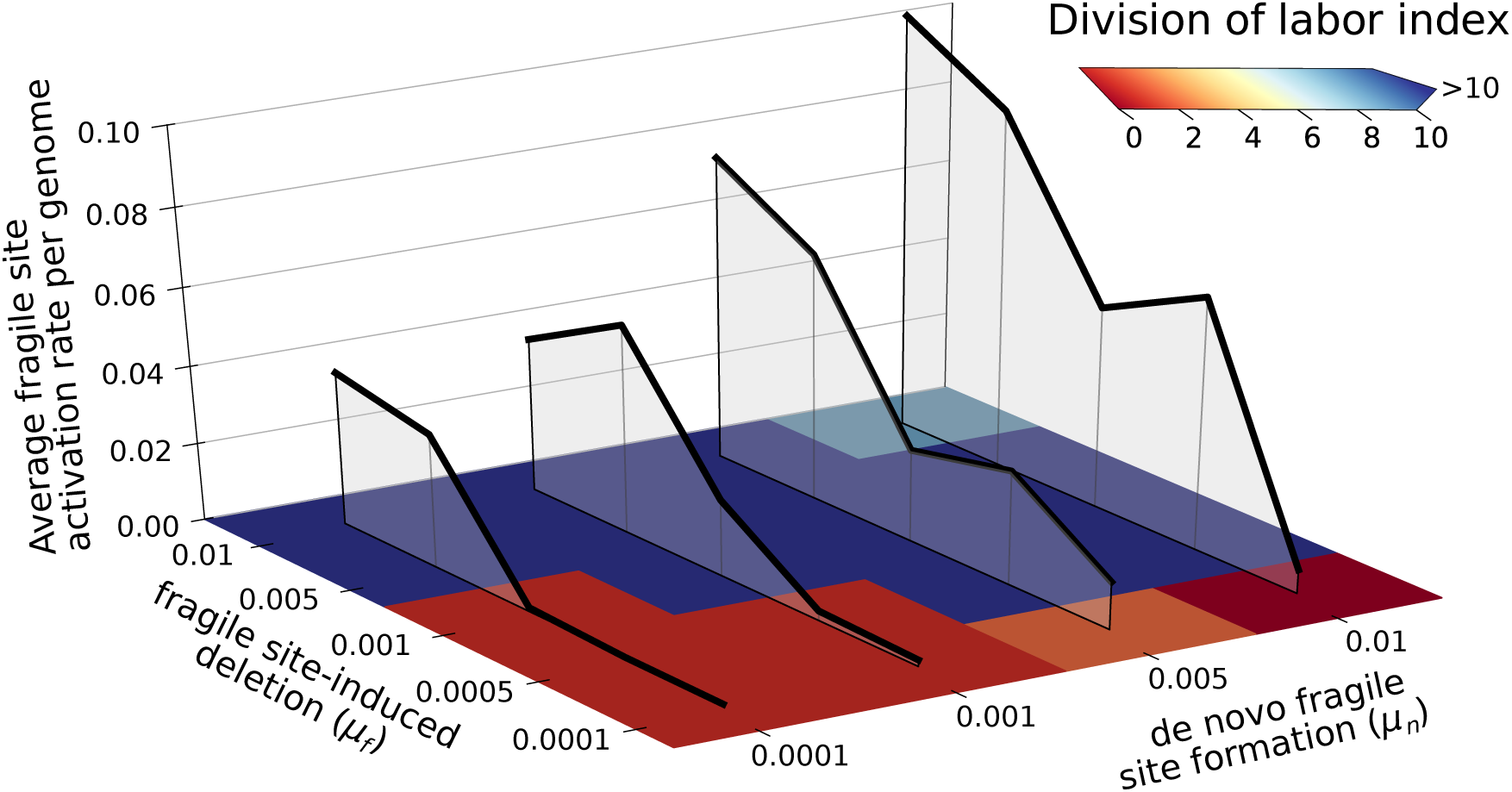
Mutation-driven division of labor and mutation rates evolve over a wide range of fragile site mutation rates. The plot shows the average genomic mutation rate (black lines) and the degree of division of labor (red to blue tiles) for a series of populations evolved under different *μ*_*n*_ (probability of *de novo* fragile site formation), and *μ*_*f*_ (fragile site-induced deletion probability). Each tile shows the difference between the median numbers of growth-promoting genes in antibiotic producers and replicating bacteria, after evolution has reached a steady state (after at least 600 growth cycles). The genomic mutation rate is calculated from the number of fragile sites *f*, as 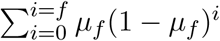.

## Discussion

We studied how an association between genome architecture and division of labor evolves, inspired by recent experimental findings in *Streptomyces* [11]. We constructed a mathematical model of *Streptomyces* multicellular development where cells have a linear genome with mutational hotspots, and experience a trade-off between replication and antibiotic production. We showed that replicating bacteria can exploit mutations to generate a new cell type that specializes in antibiotic production, thereby resolving the trade-off between these traits (Fig. 2). This mutation-driven differentiation is coordinated by an evolved genome architecture with two key features: fragile sites that destabilize the genome by causing large-scale deletions, and an over-representation of growth-promoting genes at the unstable chromosome ends. Because of this organization, mutations at fragile sites preferentially delete growth-promoting genes, leaving mutant genomes containing only genes for antibiotic production. Although these cells are unable to grow, their terminal differentiation into antibiotic-producing cells is advantageous to the colony as a whole.

*Streptomyces* genomes contain a conserved (mutationally quiet) core region located at their centromere, and a telomeric region which is prone to various forms of recombination and instability and contains much of the accessory genes [40, 41]. This leads to high within- and between-species variability at the chromosome ends [42]. Both fragile sites (including genomic inverted repeats and other recombination hotspots, associated to intra-chromosomal recombination and to mobile genetic elements) and biosynthetic clusters are unevenly distributed along the chromosome of Streptomyces [19, 41, 43], suggesting that they are positioned non randomly. Our model suggests that telomeric instabilities have evolved to coordinate the division of labor between replication and antibiotic production, explaining recent experimental results in *S. coelicolor* [11]. Thus, these instabilities may not be accidental byproducts of a linear chromosome or other aspects of unstable replication [15]; our model predicts that they are instead an evolved and functional property of *Streptomyces* genomes. Based on the robustness of our results to parameter changes, and because genome instabilities are extremely common in *Streptomyces*, we expect that other species in the genus also divide labor through this mechanism. For instance, recent results show that expression of biosynthetic pathways in S. ambofaciens is related to DNA folding [44]. Based on our model, it is tempting to speculate that fragile site-induced deletions can change DNA folding by removing parts of complementary DNA, exposing genes that were previously hidden to the transcription machinery. All these lines of evidence support the idea of the evolution of a precise genome architecture that increases the evolvability and the dispensable nature of the telomeres - providing support for the idea that the genome architecture of the telomeres in Streptomyces results from an evolutionary interplay between biosynthetic genes and fragile sites.

*Streptomyces* produce a broad diversity of metabolically expensive compounds like cellulases and chitinases that can also be used as public goods [45]. These are also expected to trade off with growth, and may, therefore, also be sensitive to division of labor through genome instability. Such examples of division of labor could alternatively be organized through differential gene expression - as is common in multicellular eukaryotes, eusocial insects and other microbes. However, an advantage of the mutation-driven division of labor presented here is that it makes social conflicts [46] impossible, because altruistic somatic cells do not possess the genetic means to reproduce autonomously and participate in social dynamics [47]. Validating these predictions will require detailed experiments in other species, as well as bioinformatic analyses of *Streptomyces* genome structures. These analyses may then inform a more detailed model of *Streptomyces* division of labor, accounting for the interplay between genome organization, gene regulation and the metabolic network, underpinning the trade-off between growth and antibiotic production.

Future models may also incorporate a more realistic mutational landscapes for biosynthetic genes and horizontal gene transfer through conjugation [48], which is known to drive novel gene acquisition and mutational hotspot formation in *Streptomyces* [19]. Moreover, exposure to some antibiotics induce DNA damage and significantly increases instability [49, 50]. Preliminary experiments have found that antibiotic production and genome instability increase in some species of *Streptomyces* during competition between colonies [51]. This suggests that the sensing of antibiotics increases mutation rates at contact points between colonies, so that mutants appear preferentially where colonies compete with one another. Investigating the consequences of this additional layer of evolutionary dynamics will be interesting in future extensions of the model.

Mathematical models indicate that the organization of genetic information along the chromosome is influenced by the mutational operators that act on it [28, 52–55]. As a consequence of this organization, mutations may be more likely to generate mutant offspring with specific characteristics, such as reduced competition with the wildtype due to low fitness [56–59], lower propensity for social conflicts [60] or accelerated re-adaptation to variable environments [61]. In prokaryotes, mutational operators that can drive functional mutagenesis include horizontal gene transfer, which drives the rapid evolution of gene content [37, 62–64], and the CRISPR-Cas system, that generates immunity to viral infections through targeted incorporation of viral genomes [65]. In an Origin of Life model, it was found that mutants could provide a benefit to the wildtype. There a germline RNA replicator evolved whose mutants were sterile but altruistically replicated the germline and protected it from parasites. [6]. Building on these earlier results, our model shows that a genome architecture can evolve to incorporate mutational hotspots, thus exploiting mutations to generate functional phenotypes and divide labor.

Beyond *Streptomyces*, mutation-driven division of labor occurs in the genome of many ciliates, where functional genes must be carefully excised from a transposon-riddled genomic background before being transcribed [66, 67]. Programmed DNA elimination in somatic cells is common in multicellular eukaryotes [68], and targeted recombination is essential for the functioning of the adaptive immune system in vertebrates [69]. These examples highlight the ubiquity of functional mutagenesis across the tree of life. Our model may therefore help understand the evolutionary origin of division of labor through functional mutagenesis more broadly.

### Conclusions

Our model shows that a bacterial colony can divide labor between replication and antibiotic production by evolving a genome architecture that physically segregates genes for these different functions, and which allows them to be dissociated due to mutations at fragile sites. This gives rise to an effective germ-soma division driven by mutation. One class of cells specializes on reproduction, while the other class, comprising the mutants, focuses on costly antibiotic production. In this system, which replicates dynamics observed in *S. coelicolor*, mutant cells function as soma by enhancing colony fitness, despite being sterile.

## Methods

The model is a lattice-based stochastic simulation system. We consider a population of bacteria that can replicate and produce antibiotics. The model is inspired by the life cycle of *Streptomyces coelicolor*, and focuses on the hyphal growth phase, when colonies develop and compete by producing antibiotics [2]. We therefore model the eco-evolutionary dynamics occurring during several colony growth cycles, each of fixed duration *τ*_*s*_. At the beginning of each cycle, spores germinate and colonies begin to form through bacterial replication. If antibiotics are produced, they are deposited around the producing bacterium, forming a halo that protects the colony and “reserves” space to replicate into. Bacteria die if they come into contact with antibiotics to which they are sensitive, and therefore bacteria of a colony cannot invade into the antibiotic halo of another one - if they are genetically different. At the end of the cycle, corresponding to the sporulation phase in *Streptomyces*, a small random sample *ξ* of the population is selected to form spores. These spores seed the next cycle, and other bacteria and antibiotics are removed from the lattice. Spores are deposited at the same location where the bacterium lived (we do not shuffle the location of the spores), unless explicitly stated.

We model bacteria and antibiotics on two separate lattices Λ_1_ and Λ_2_. Both lattices have size *L × L* and toroidal boundaries to avoid edge effects. Every site of Λ_1_ can either be occupied by one bacterium, or be empty. Every site of Λ_2_ can be occupied by multiple antibiotics. Each bacterium possesses a genome which determines their replication rate, as well as the rate and type of antibiotics it produces and is resistant to. The genome consists of a linear sequence of three types of genetic elements (so called beads on a string model [28, 29]). We consider two gene types - growth-promoting genes and antibiotic genes. The third type of genomic element is a fragile genomic site - a hotspot for recombination. Growth-promoting genes increase replication rate and inhibit antibiotic production, in accordance with *Streptomyces* growth being favored over secondary metabolism. Antibiotic genes encode both the toxin and its resistance ([70]). Antibiotic type is encoded in the genetic sequence of the antibiotic gene. This sequence is modelled as a bit-string of fixed length *ν*, which can be mutated to encode different antibiotics. Each bacterium can have multiple growth-promoting and antibiotic genes, as well as multiple fragile sites.

### Replication

Replication rate depends on the intrinsic replication rate function *G* and on the resistance *R* of the bacterium to antibiotics in the spatial location in which it lives:

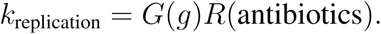

The intrinsic replication rate *G*(*g*) is an increasing, and saturating, function of the number of growth-promoting genes *g*:

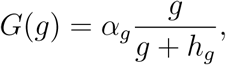

with *α*_*g*_ the maximum replication rate, and *h*_*g*_ the number of growth-promoting genes producing half maximum replication rate. Antibiotic resistance *R* depends on whether a genome has at least an antibiotic gene sufficiently similar to the antibiotics present on Λ_2_ at the corresponding location of the bacterium, where similarity is determined from the bit-strings of the antibiotics and the antibiotic genes. For each antibiotic *a* on Λ_2_, the Hamming Distance *D* (the number of different bits) between the antibiotic and the gene with the minimum distance from *a* is calculated. All these minimum distances are summed into a susceptibility score *S* =∑*D*, and the overall resistance is a decreasing function of *S*:

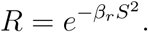

Each bacterium is resistant to the antibiotics it produces, because for each antibiotic *D* = 0, and thus *R* = 1. Moreover, the Gaussian function ensures that small mutations in antibiotic types do not decrease resistance too rapidly. Replication occurs when bacteria are adjacent to an empty site on Λ_1_. The probability of replication for each competing bacterium *i*, is *k*_replication_(*i*)*/η*, with *η* the neighborhood size (we use the Moore neighborhood throughout this study, thus *η* = 8). Upon replication, the new bacterium inherits the genome of its parent, with possible mutations.

### Mutations

Mutations occur during replication, and can expand or shrink genomes, and diversify antibiotics.

#### Duplications and deletions

Duplication and deletion of genes and fragile sites occur with equal per-gene probability *μ*_*d*_. When a gene or a fragile site is duplicated, the gene copy is inserted at a random genomic location. This ensures that gene clustering is not a trivial consequence of neutral mutational dynamics, and must instead be selected upon to evolve.

#### Fragile site large chromosome deletions

Fragile sites are the cause of genome instability in the model. We let fragile sites cause large-scale chromosomal mutations with a per-fragile site probability *μ*_*f*_. We take into account that large-scale mutations in *Streptomyces* preferentially disrupt telomeric regions [15, 16, 18, 20] by letting fragile site-induced mutations delete the entire telomeric region offstream of the genomic location of the fragile site (see Fig. 1c).

#### Mutations of the antibiotic genes

The antibiotic genes consist of a genetic sequence modelled as a bit-string of length *ν*. Mutations flip bits with a uniform per bit (per antibiotic gene) probability *μ*_*a*_. This changes the antibiotic type, and thus the antibiotic repertoire of the bacterium.

#### Influx of new fragile sites

Fragile genomic sites, such as inverted repeats or transposable elements are common in the genome of *Streptomyces*. Because they are easily copied (or translocated) we assume that they can also be spontaneously generated with a small probability *μ*_*n*_ (independent of genome size). The new fragile site is inserted at a random location in the genome.

### Antibiotic production, in a trade-off with replication

Antibiotic production rate is modeled as an increasing function *A* of the number of antibiotic genes *a* a bacterium has. At the same time, production is strongly inhibited by growth- promoting genes - a function *I*(*g*) with *g* the number of growth-promoting genes in the genome, in accordance with a likely trade-off between growth and secondary metabolism [11]. Antibiotic production rate, per time step, is the function:

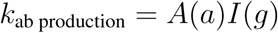

with

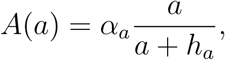

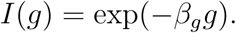

According to this function, the trade-off becomes rapidly steeper with larger *β*_*g*_. For simplicity, we assume that antibiotics are deposited at a random location within a circle of radius *r*_*a*_ around the producing bacterium. Moreover, we do not take into account concentrations of antibiotics and only model presence/absence of an antibiotic type in a spatial location. The probability of producing an antibiotic, per site within the circle, per unit-time is equal to *k*_ab production_. If a bacterium has multiple antibiotic genes, the antibiotic deposited is chosen randomly and uniformly among them.

### Death

Death can occur when a bacterium is sensitive to an antibiotic located at the same site as the bacterium itself. The probability of a bacterium dying is calculated as 1 − *R*, where *R* is the bacterial resistance to the antibiotic defined above. When a bacterium dies, it is removed from the lattice, leaving behind an empty site.

### Movement

Bacteria have a small probability *p*_mov_ of moving, if there is an empty site adjacent to them. This speeds up colony expansion and competition between colonies, and avoid strong grid effects that could make spatial patterns too rigid.

### Initial conditions and updating of the dynamics

Unless differently specified, at time *t* = 0 a small population of spores is seeded on the lattice. The initial spores have a small genome of length 10, i.e. a random sequence of growth-promoting genes and antibiotic genes (of random types), but no fragile sites. The lattice is updated asynchronously: over one time step, each lattice site is updated in random order.

The source code is written in c, and uses the CASH libraries [71]. Analysis and plotting custom software is written in python. Genome schematics in Fig. 3b are partly drawn with dnaplotlib [72]. The source code and analysis scripts are available at [73].

## Supplementary Material

### S1 The evolutionary dynamics of genome composition

For clarity, in the main text we showed only one example of the evolutionary dynamics. Fig. SF1 shows that qualitatively the same evolutionary trends are followed when the experiment is repeated. The figure also shows that population size at the end of each growth cycle remains approximately constant at about 60000 individuals over evolutionary time.

The mutation-driven division of labor described in the main text is robustly driving the eco-evolutionary dynamics also when we run the system for a very long time (2000 growth cycles), see Fig. SF2. Interestingly, we also observe large fluctuations in the number of antibiotic genes.

**Figure SF1:**
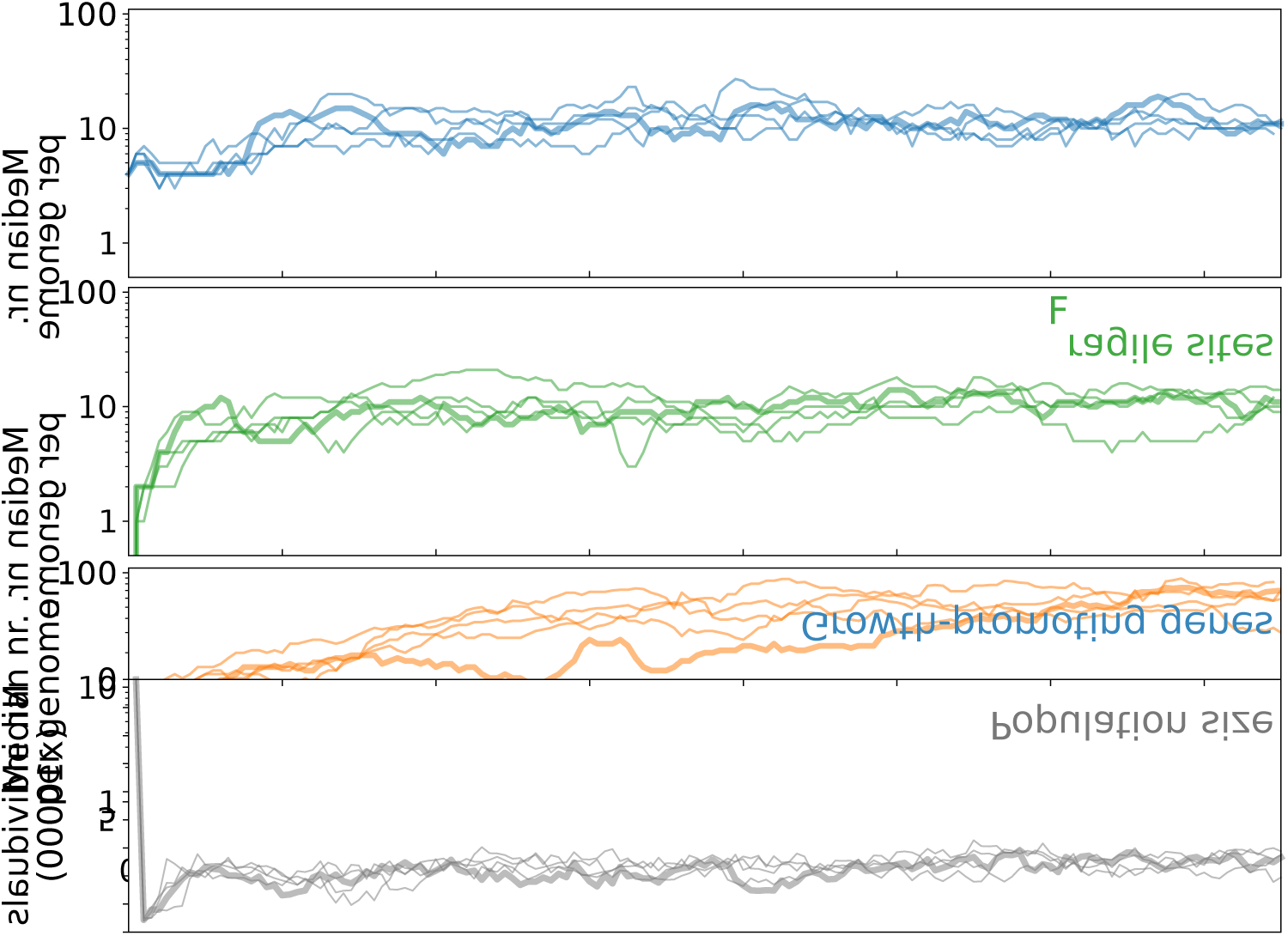
Evolutionary dynamics of population size and gene content, in five independent evolutionary runs. The run used in the main text is indicated with a thicker line. All the runs are initialized with a population with genome: `5’-FAAFAAFAAF-3’`, where F is a growth-promoting gene and A is an antibiotic gene, with each antibiotic gene encoding a different antibiotic type. Top plot shows the total population size at the end of each growth cycle. The other plots show the median number of each type of genetic element, calculated from all genomes in the population at the end of every growth cycle. Growth cycle duration *τ*_*s*_ = 2500 time steps.

**Figure SF2:**
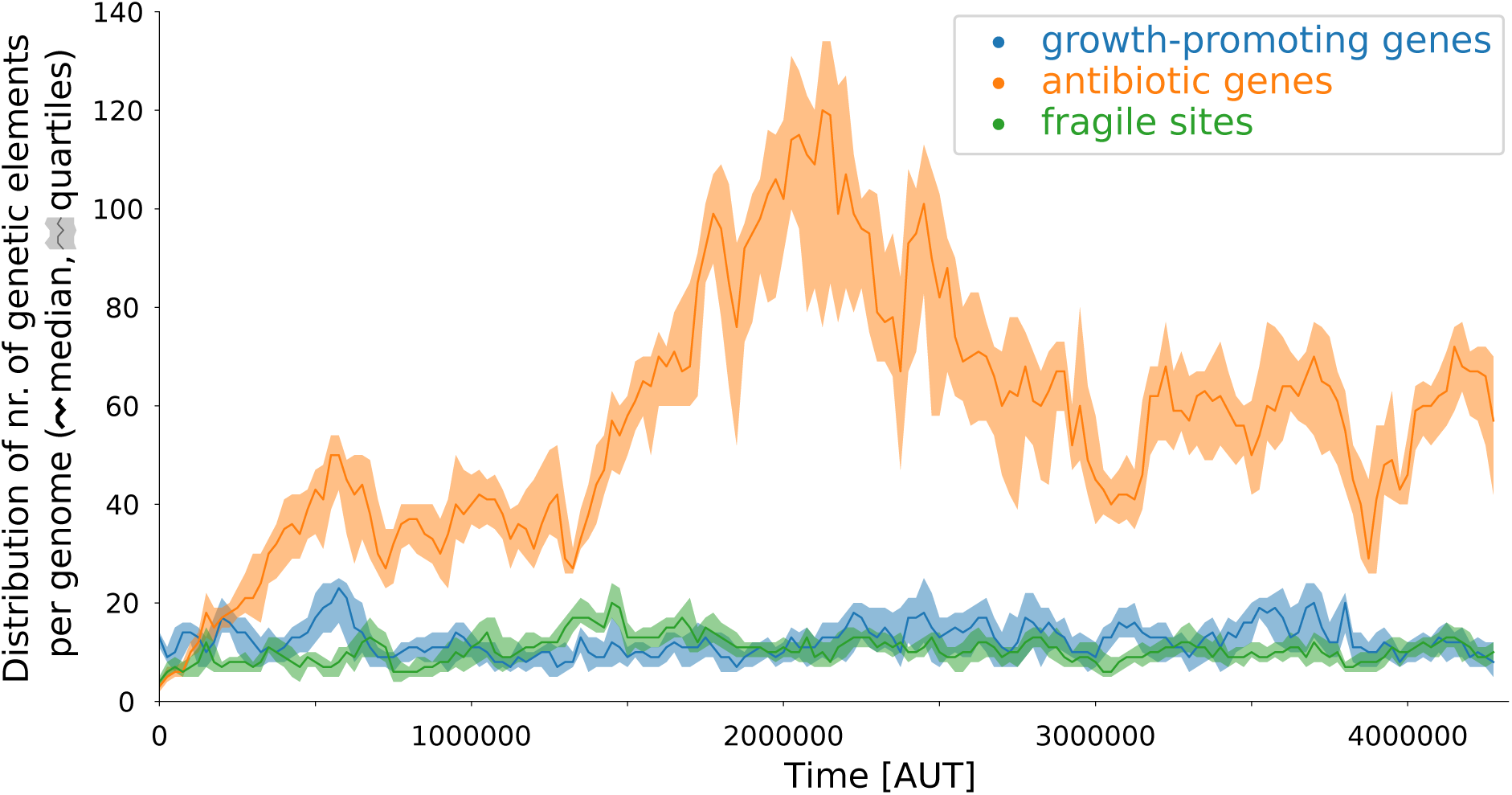
Very long-term evolutionary dynamics of genome composition. The median and quartile values of the number of each type of genetic element are calculated from all genomes in the population, at the end of every growth cycle. All parameters are the same as in Fig. 2a. The system was initialized with a small population of randomly generated genomes of length = 20 consisting, on average, of 65% growth-promoting genes, 15% antibiotic and 20% fragile sites. Antibiotic type was also randomly generated. Growth cycle duration *τ*_*s*_ = 2500 time steps.

### S2 Snapshot of the eco-evolutionary dynamics within one growth cycle

Fig. SF3 shows successive snapshots of the lattice, over the course of a growth cycle that lasts 2500 time steps. Starting from spores, colonies expand and produce antibiotics. Antibiotic-producing mutants form a dotted pattern over the colony. For each time point, the top pane shows colonies and antibiotics, the bottom pane shows antibiotic potential in non-producing cells (higher with darker blue) and antibiotic-producing cells (red).

**Figure SF3:**
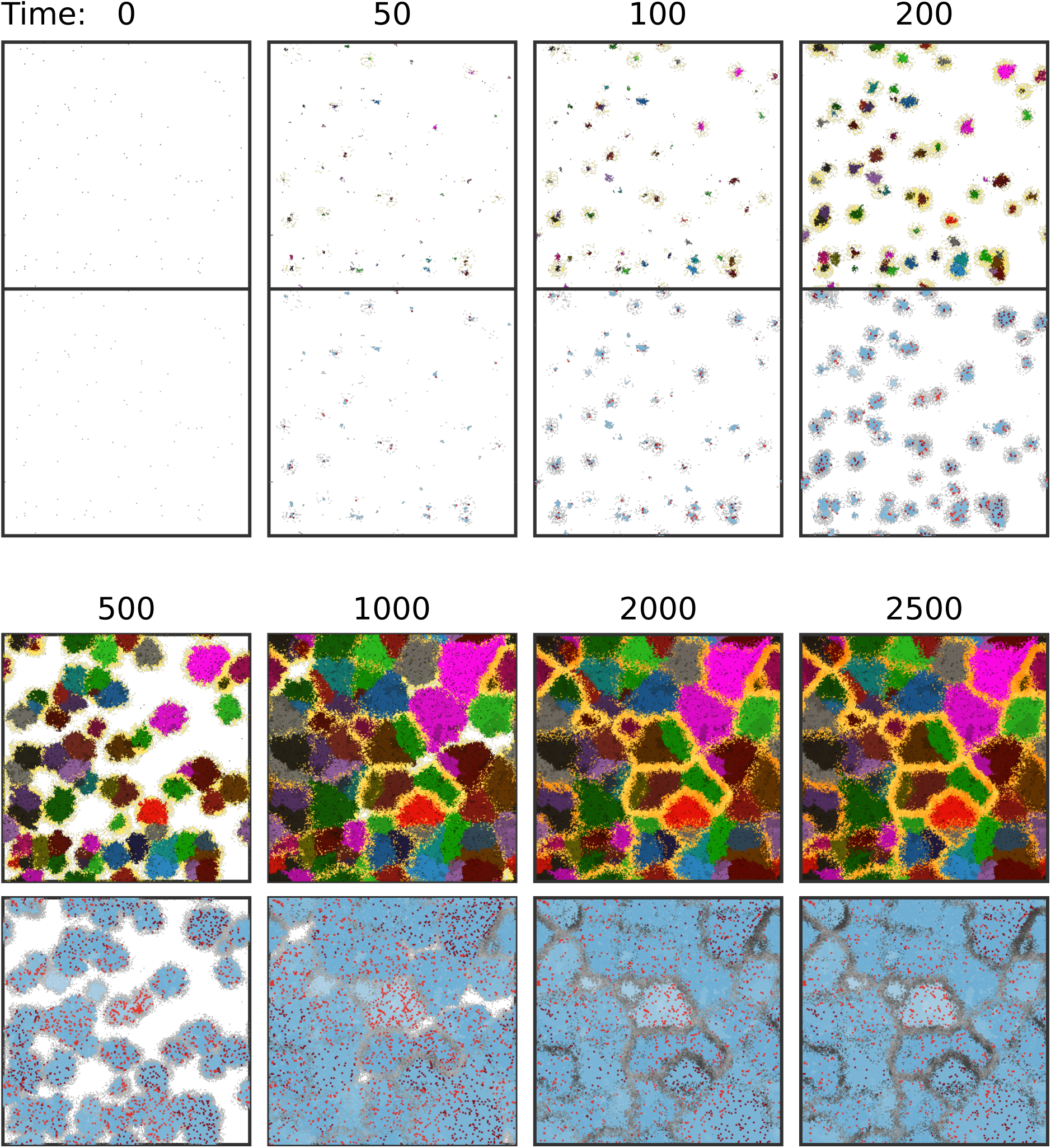
The eco-evolutionary dynamics during a growth cycle. For each time point, top pane: different colors represent colonies arising from single spores, darker shades of yellow indicate that more antibiotics are present; bottom pane: darker shades of blue indicates larger number of antibiotic genes in non-producing (and weakly producing) cells, red dots indicate antibiotic-producing bacteria (brighter red corresponds to higher production), gray corresponds to antibiotics. In both panes, white is background.

### S3 The mild fitness costs of dividing labor through mutation

We perform several competition experiments between wildtype bacteria that divide labor through mutations, and “artificial” generalists whose replication rate and antibiotic production rate are drawn from a broad spectrum of values. We chose three independently evolved bacteria, and repeat each competition experiment four times. We initialize the system with a mix of the two genotypes, and we run the simulation with mutation rates *μ*_*d*_, *μ*_*n*_, *μ*_*a*_ set to zero (so that no further evolution happens), and with the probability of fragile-site deletion *μ*_*f*_ set to default values (so that division of labor can occur). After 15 growth cycles, the winner of the competition is determined. Fig. SF4 shows the sum of how many times the artificial generalist (+1) or the wildtype (−1) wins, for each combination of the generalist’s growth and antibiotic production rate. A draw occurs in some cases (0), when both genotypes are present in similar numbers after 15 cycles. The figure also shows the replication rate and antibiotic production rate of the wildtype species we used (yellow bar). For these, replication rate was calculated from the number of growth-promoting genes in their genome (see Methods), while average and standard deviation of per-capita antibiotic-production rate was obtained from a simulation which was initialized with only this bacterium, and all mutation rates except *μ*_*f*_ were set to zero (to avoid evolution). The three wildtype genomes are as follow (A: antibiotic genes; F: growth-promoting genes; B: fragile sites):

~~~
>Wildtype Genome 1: nr. F=14, nr. A=111, nr. B=15 AABAAABABAAAAAAAAAAAABAAAAAABAAAAABAAAAAAAABAAAAAABAABABAABA AAABAABAFAAAAAAAABFAAAAAAAFAAAAFAAAAFFAAFAFAAAAAFAAAAAAAAAAA AABAAFAAAFAFAAFAAFAA
>Wildtype Genome 2: nr. F=21, nr. A=61, nr. B=10 AAAAAAAAAABAAABAAAAAABAABABABBBAAFABAAAAAAFFAAAAAAAFAAAFAFFA AAFFAFFFBAFAAFFAFAFFAFAAAAAFAAAF
>Wildtype Genome 3: nr. F=15, nr. A=71, nr. B=14 AAAABAABAAAAAAAAABAAABAAAAAFAAABBAAABAAAABABAABAAAAAFFFBAABA FABAFAAAAAFFAFAAFAFAAAABAFAAAAFAFAAAFAAA
~~~

**Figure SF4:**
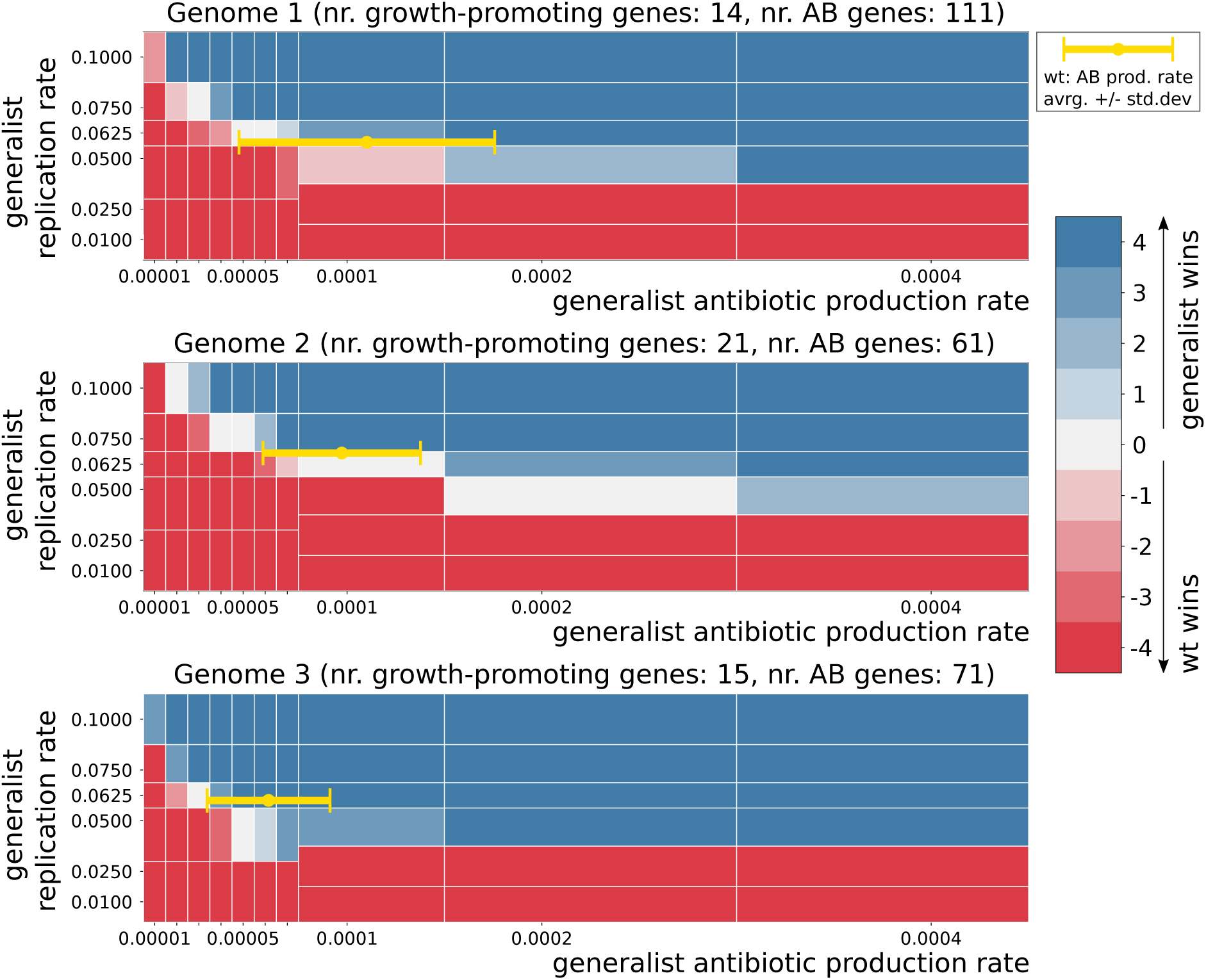
Competition experiments between artificial generalists and three evolved wild type species. The plot shows the number of times the generalist or wildtype wins, given different values of growth rate and antibiotic production rate of the generalist. Blue: majority generalist wins. Red: majority wildtype wins. Grey: majority draws or equal number of wins.

### S4 The evolution of division of labor depends on the trade-off between antibiotic production and replication - the effect of *β*_*g*_ and *h*_*g*_

#### Larger *β*_*g*_ favors division of labor

Antibiotic production rate *k*_ab_ is inversely related to the number of growth-promoting genes via the equation *k*_ab_ = *A* exp(−*β*_*g*_*g*) (see Methods) - effectively imposing a trade-off between replication and antibiotic production. The strength of the trade-off can be tuned by the parameter *β*_*g*_, which in the main text is set to *β*_*g*_ = 1 throughout. The closer *β*_*g*_ is to zero, the more antibiotic production becomes independent of the number of growth genes. Very small *β*_*g*_ corresponds to an unrealistic situation where bacteria have arbitrary energy to maximize both replication and antibiotic production. We expect that no division of labor evolves in this case. Indeed, Fig. SF5 shows that for *β*_*g*_ *<* 0.5, division of labor does not evolve. The figure also shows that for *β*_*g*_ = 0.5 division of labor occasionally evolves - but it is evolutionarily unstable (not shown). Interestingly, the plots also show that changing *β*_*g*_ does not affect the steady state composition of the genome. As shown below and in the main text, parameters controlling mutations and growth have a larger influence on genome composition.

#### Larger *h*_*g*_ favors division of labor

Replication rate depends on the number of growth genes in the model. The parameter *h*_*g*_ sets the number of growth-promoting genes *g* that result in half-maximum growth rate, i.e. increasing *h*_*g*_ results in a lower growth rate for a fixed value of *g*. Trade-off strength can be increased with larger *h*_*g*_, because larger *g* will be required for growth, which results in a decrease in antibiotic production rate. Indeed, Fig. SF6 shows division of labor evolves for *h*_*g*_ *≥* 6. For *h*_*g*_ = 4 division of labor evolves, but we observe that it is evolutionarily unstable.

**Figure SF5:**
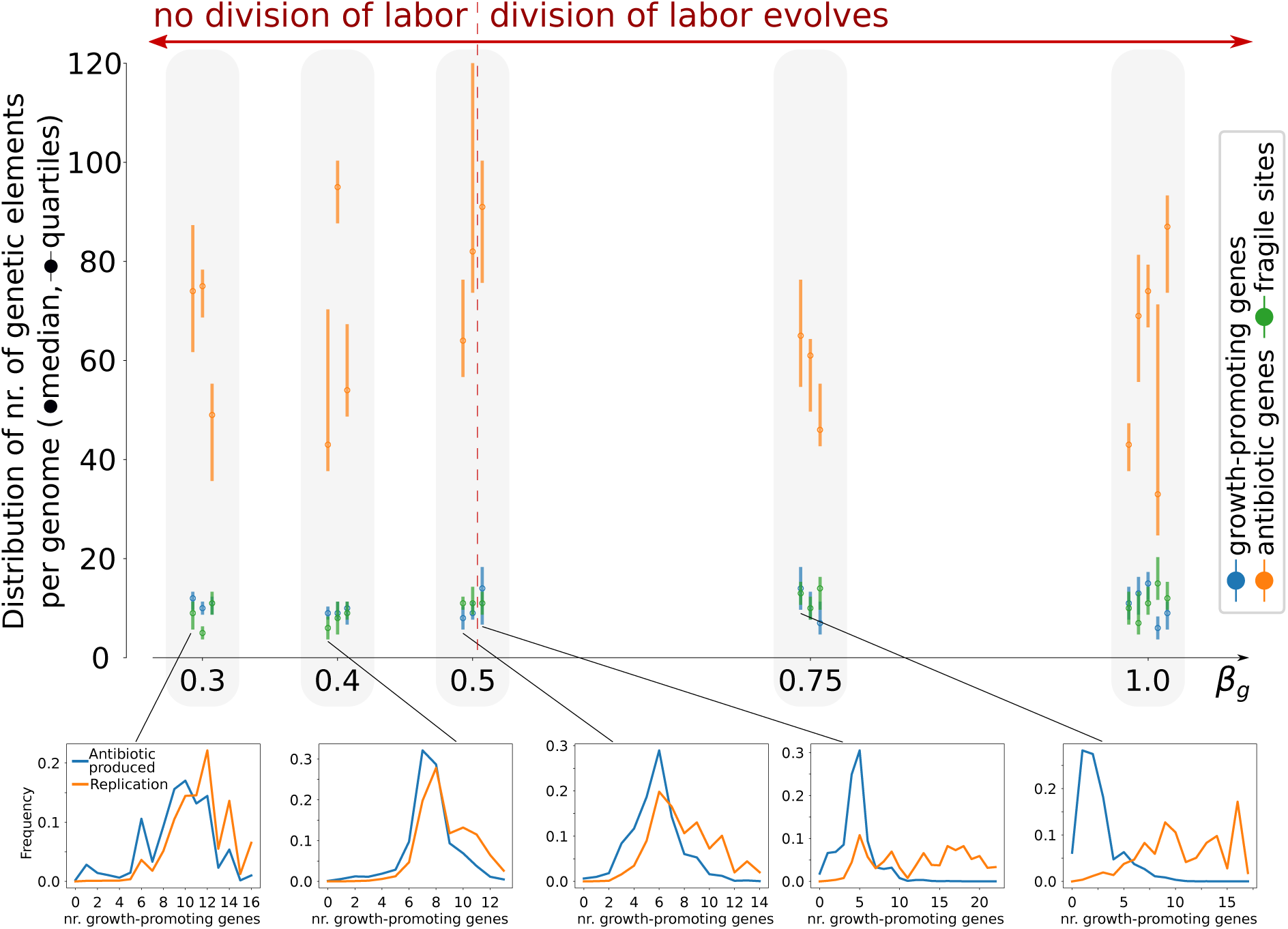
Division of labor evolves when inhibition of antibiotic production is sufficiently strong with a large nr. of growth-promoting genes (thus making the trade-off between replication and antibiotic production stronger). Data is collected from the entire population, for one growth cycle after long-term evolution (*>* 1000 generations) from a series of simulations with different values of *β*_*g*_ (three replicas for each *β*_*g*_; all other parameters are identical to those in the caption of Fig. 1). The top pane shows the distribution of each genetic element in the genomes. Bottom panes show the frequency of antibiotic producers (blue) and replicating individuals (orange) as a function of the nr. of growth-promoting genes, for the replica indicated in the figure. A larger difference between the two curves indicates division of labor, because the two tasks are carried by genetic distinct individuals in the same colony.

**Figure SF6:**
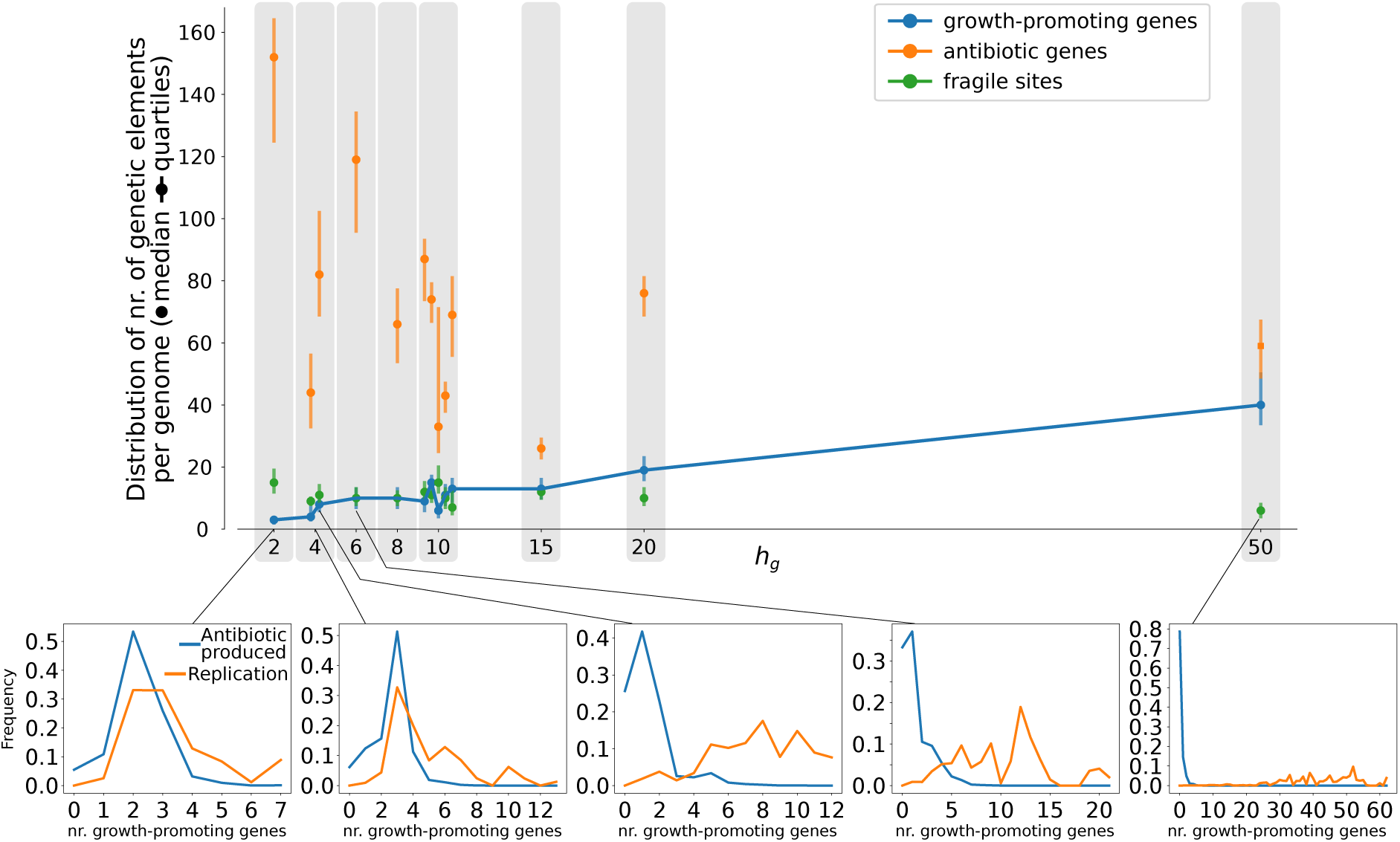
Division of labor evolves when the nr. of growth genes required for half-maximum growth is sufficiently large, making the trade-off between replication and antibiotic production stronger. Data is collected from the entire population, for one growth cycle after long-term evolution (*>* 1000 generations) from a series of simulations with different values of *h*_*g*_ (all other parameters are identical to those in the caption of Fig. 1). The top pane shows the distribution of each genetic element in the genomes. Bottom panes show the frequency of antibiotic producers (blue) and replicating individuals (orange) as a function of the nr. of growth promoting genes, for the replica indicated in the figure. A larger difference between the two curves indicates division of labor, because the two tasks are carried by genetic distinct individuals in the same colony.

### S5 A scaling relationship between growth and antibiotic production controls trade-off strength

The growth function can be written as: 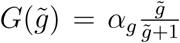, where 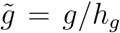, showing that growth rate as a function of *g* is measured in units of (1*/h*_*g*_). Similarly, antibiotic production (as a function of *g*) can be written as 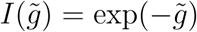, where 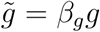, showing that antibiotic production rate as a function of growth-promoting genes is measured in units of *β*_*g*_. If we set *β*_*g*_ = *K/h*_*g*_, we can change antibiotic production rate and growth rate maintaining the ratio between the two functions. Since changing either parameter changes trade-off strength (see Suppl. Section S4), but setting *β*_*g*_ = *K/h*_*g*_ does not, we conclude that the product *β*_*g*_*h*_*g*_ scales trade-off strength. Indeed, Fig. SF7 division of labor evolves when we change the values of *β*_*g*_ and *h*_*g*_ maintaining their product to *K* = 10.

**Figure SF7:**
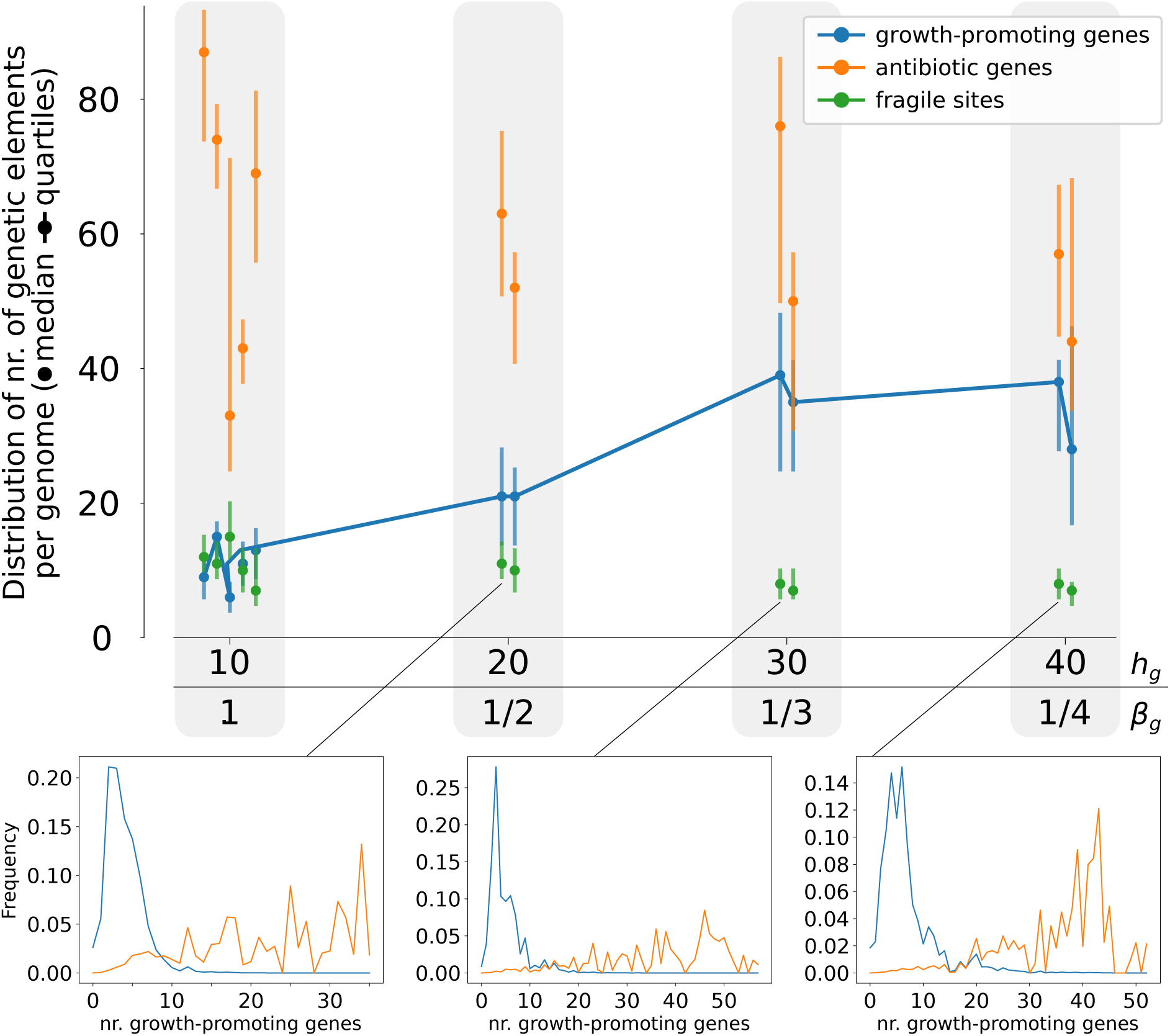
Trade-off strength is kept constant - and division of labor can evolve - by scaling the nr. of growth genes required for half-maximum growth with the nr. of growth genes required for inhibition of antibiotic production. Data is collected from the entire population, for one growth cycle after long-term evolution (*>* 1000 generations) from a series of simulations with different values of *h*_*g*_ and *β*_*g*_ such that *h*_*g*_*β*_*g*_ = 10 (all other parameters are identical to those in the caption of Fig. 1). The top pane shows the distribution of each genetic element in the genomes.. Bottom panes show the frequency of antibiotic producers (blue) and replicating individuals (orange) as a function of the nr. of growth promoting genes, for the replica indicated in the figure. A larger difference between the two curves indicates division of labor, because the two tasks are carried by genetic distinct individuals in the same colony.

### S6 Weaker trade-off and lower overall antibiotic production enable division of labor

We show that trade-off strength is not the only determinant of the evolution of division of labor. The overall antibiotic production rate is controlled by the parameter *α*_*a*_. When *α*_*a*_ is smaller, fewer antibiotics are produced by bacteria, even when they have few growth-promoting genes. Thus, larger *α*_*a*_ might be expected to favor division of labor. Surprisingly, we show in Fig. SF8 that smaller values of *α*_*a*_ enable division of labor when the trade-off is shallower (i.e. when *β*_*g*_*h*_*g*_ is small). We set *h*_*g*_ = 10 - as in main text, and *β*_*g*_ = 0.5, resulting in *β*_*g*_*h*_*g*_ = 5, which makes the trade-off too shallow for division of labor to robustly evolve. We further set *α*_*a*_ = 0.075, which is more than an order of magnitude smaller than default value (the specific value of *α*_*a*_ is chosen so that the antibiotic production rate for an intermediate nr. of growth promoting genes (*g* = 5) is the same as that obtained with default values. In formulas, *I*(*α*_*a*_ = 0.075, *β*_*g*_ = −0.5, *g* = 5) *≈ I*(*α*_*a*_ = 1, *β*_*g*_ = −1, *g* = 5)). This suggests that division of labor can evolve under mild conditions: it suffices that fast replicating individuals cannot make enough antibiotics for the colony, and is thus a likely outcome of the evolutionary dynamics of antibiotic-producing microbes.

**Figure SF8:**
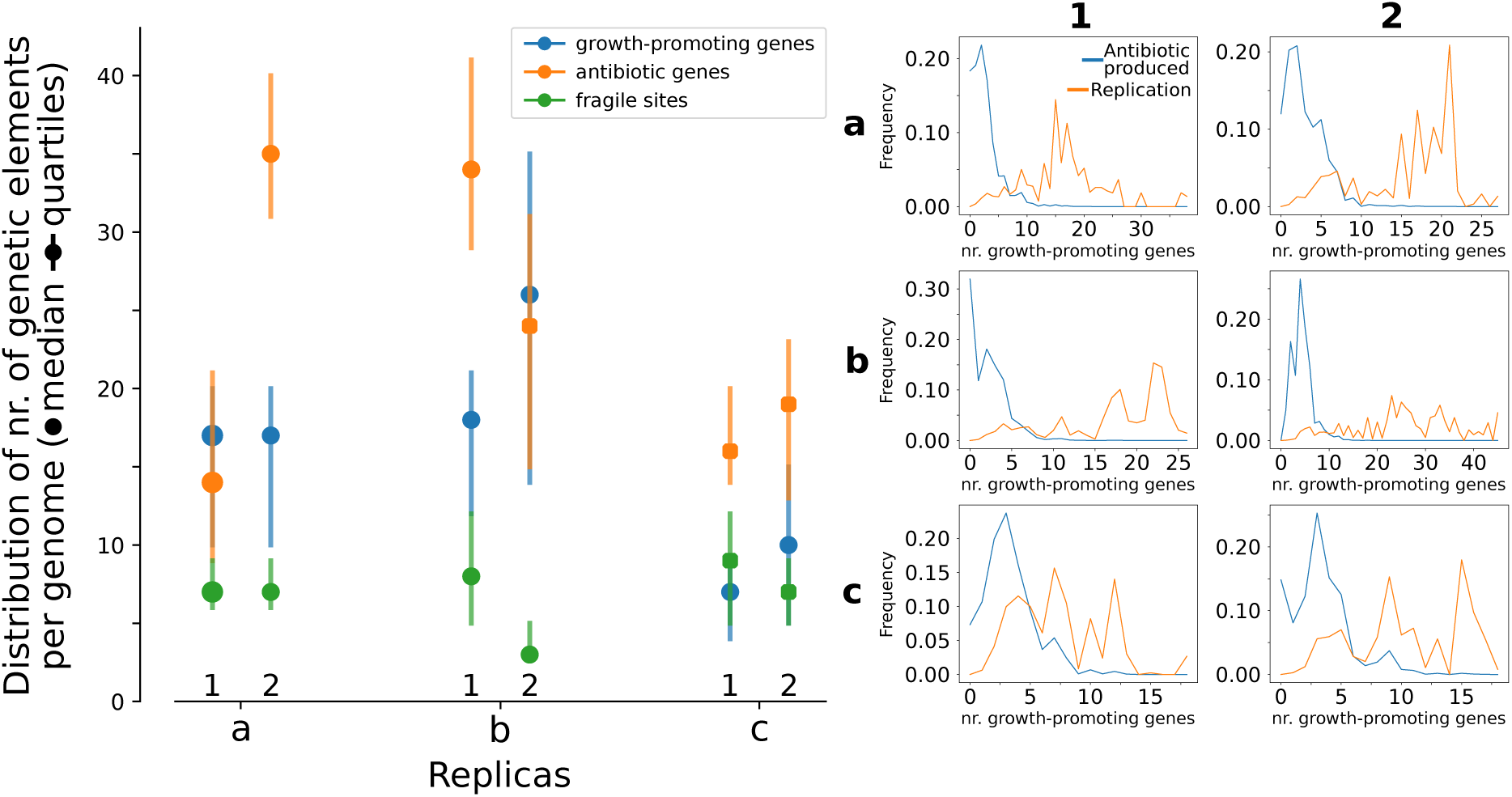
Weak trade-off and low antibiotic production allow for evolution of division of labor. Three independent simulations (a,b,c) are run with parameters *h*_*g*_ = 10 and *β*_*g*_ = 0.5 and *α*_*a*_ = 0.075 (all other parameters are identical to those in the caption of Fig. 1). Data is collected from the entire population, for one growth cycle after long-term evolution from two data points (at generation nr. 1600 and 2000). The left pane shows the distribution of each genetic element in the genomes after long-term evolution. Right panes show the frequency of antibiotic producers (blue) and replicating individuals (orange) as a function of the nr. of growth promoting genes, for the replicas indicated in the figure. A larger difference between the two curves indicates division of labor, because the two tasks are carried by genetic distinct individuals in the same colony.

### S7 The fraction of mutants during colony development

As shown in the main text, colonies begin their growth cycle clonally, and diversify through mutations. Mutants that overproduce antibiotics show massive deletions in their telomeres and lack growth promoting genes. We select mutants that deleted at least 3*/*4 of their growth genes and have at least 1 antibiotic gene as a proxy for mutants that hyperproduce antibiotics (see main text Fig. 2d). Fig. SF9 shows how the fractions of these mutants changes during colony development. For colonies at an early time stage (i.e. at 40 growth cycles), mutation-driven division of labor has not evolved yet, and therefore mutations are largely deleterious and over a growth cycle mutants are outcompeted by the wild-type. At later stages, after division of labor has evolved, the fraction of mutants is much larger than for earlier cases reaching up to 7% of the population at the beginning of the growth cycle, and stabilizing above 2% at the end.

**Figure SF9:**
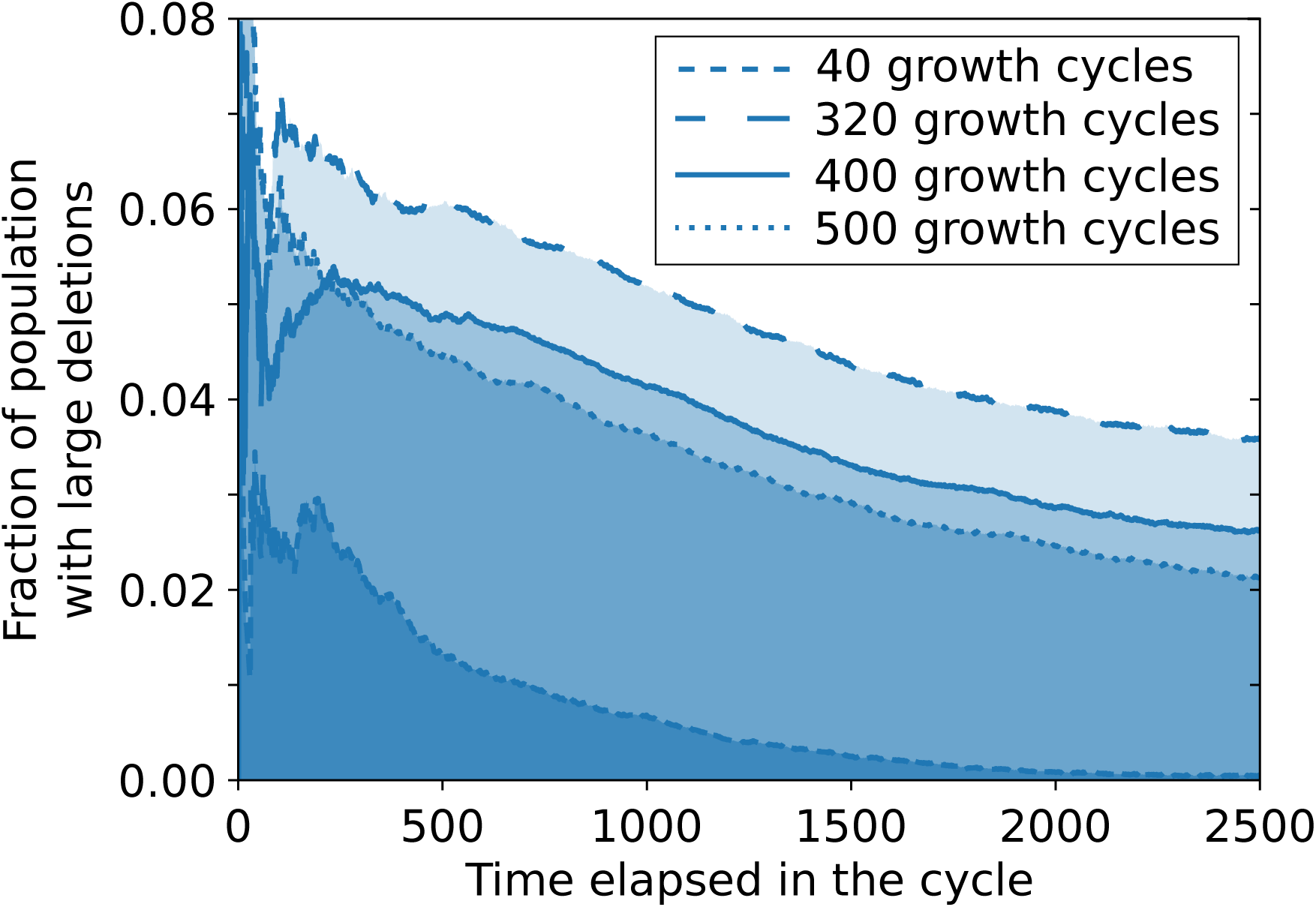
The fraction of mutants during colony development (i.e. over one growth cycle of 2500 time steps), for different time points. At growth cycle 40, division of labor has not evolved yet. At growth cycle 320, 400 and 500 division of labor has evolved and mutants are stably maintained throughout the growth cycle.

### S8 Genome architecture: growth-promoting genes

Fig. SF10 shows that the evolved genome architecture compartmentalizes growth-promoting genes to the telomeric region of the bacterial chromosome. While in the main text we presented one example colony, here we show that the evolved genome architecture is similar in the entire population. We extract the genome of all the individuals at the end of one growth cycle, after long-term evolution (more than 1000 growth cycles), from the simulation shown in Fig. SF2. Because genome size is highly variable in the population, we normalize the position of each gene by the genome length, so that position 0 corresponds to the most centromeric region, and 1 corresponds to the most telomeric region. We then generate a 2D histogram that correlates the position of each growth-promoting gene with the size of the genome. To avoid correlation between genome size and number of growth-promoting genes, we normalize the contribution of each gene by the size of the genome it comes from.

**Figure SF10:**
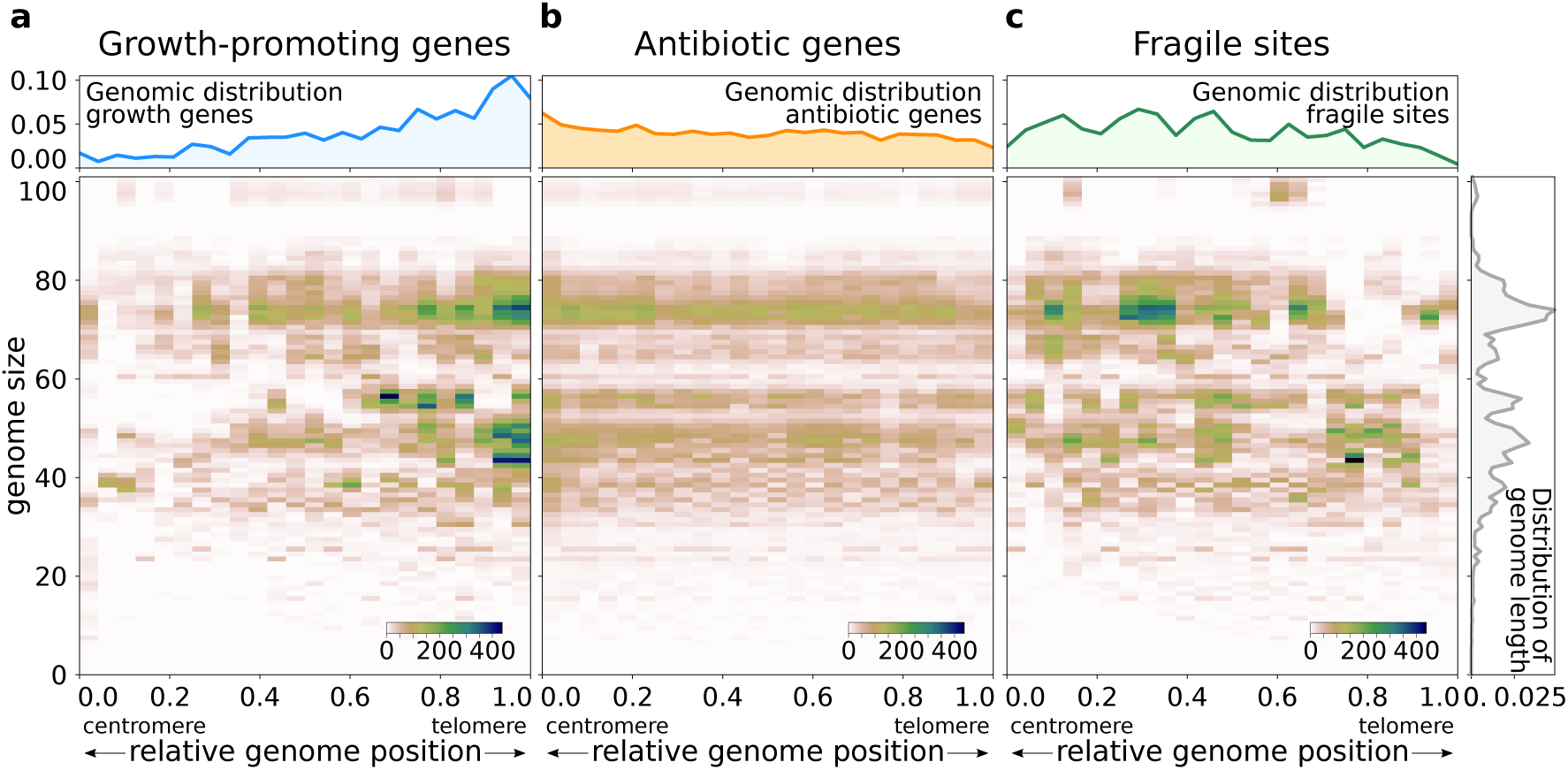
The evolved genome architecture: growth promoting genes are compartmentalized to the telomeric side of fragile sites, so that fragile-site deletions growth genes in block. Data from the last time step of the simulation shown in Fig. SF2.

In Fig. SF11, we also show the genome architecture extracted from the populations at the last time point of the five simulations shown in Fig. SF1.

**Figure SF11:**
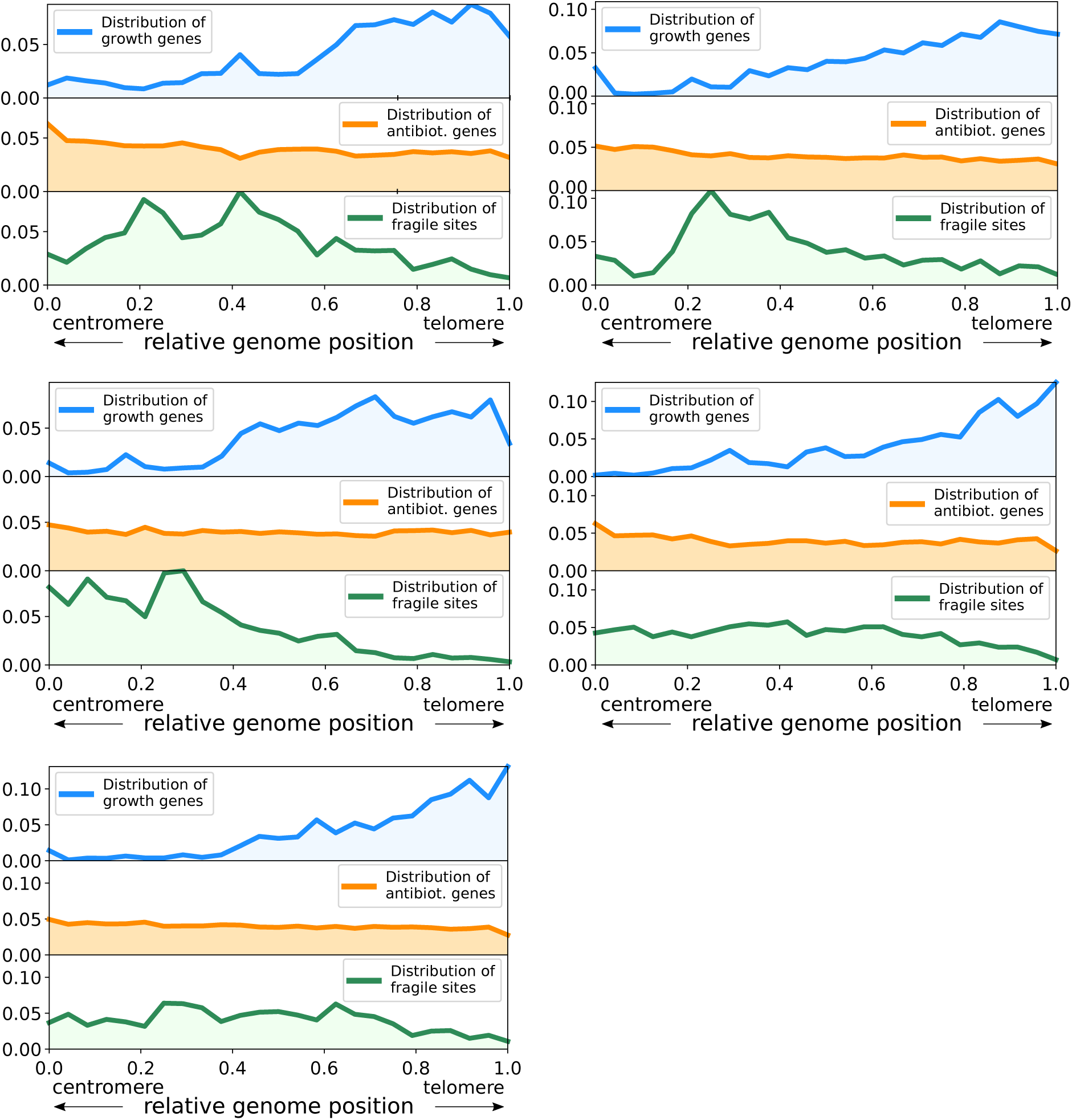
The evolved genome architecture is a very robust outcome of the evolutionary dynamics. Each pane corresponds to (and gathers data from the last time step of) one of the five simulations shown in Fig. SF1.

### S9 Competition between wildtype and genomes evolved with shuffling

Bacteria that evolved division of labor under default settings, always win in the competition with genomes evolved with genome shuffling (main text Fig. 4). We choose three wildtype genomes (see Supplementary Section S3 for genome sequence) and two genomes that evolved with sequence shuffling, and perform all pairwise comparisons (10 times) between the two types following the same protocol as in Supplementary Section S3. In all cases, the wildtype genome wins over the one evolved with shuffling.

The sequences of the genomes evolved with shuffling are:

~~~
>Shuffling Genome 1: nr. F=5, nr. A=23, nr. B=1 ABAAAAAAAAAFFAAAAAAAAAAAFFFAA
>Shuffling Genome 2: nr. F=7, nr. A=28, nr. B=1 AFAAAAFAAAAABAAAAAFAAFFAAAAAAAAFAAFA
~~~

### S10 Evolution of genome architecture with an additional gene type

Fig. SF12 shows that an additional gene type, assumed to be essential for survival in at least *n*_*h*_ = 10 copies, evolves to be correctly partitioned towards the centromere of the chromosome. With this evolved genome organization, these genes do not compromise division of labor because they are placed to the 5’ of fragile sites, thus ensuring survival of both a wildtype bacterium with a complete genome and of an antibiotic-producing mutant arising through fragile site deletion.

**Figure SF12:**
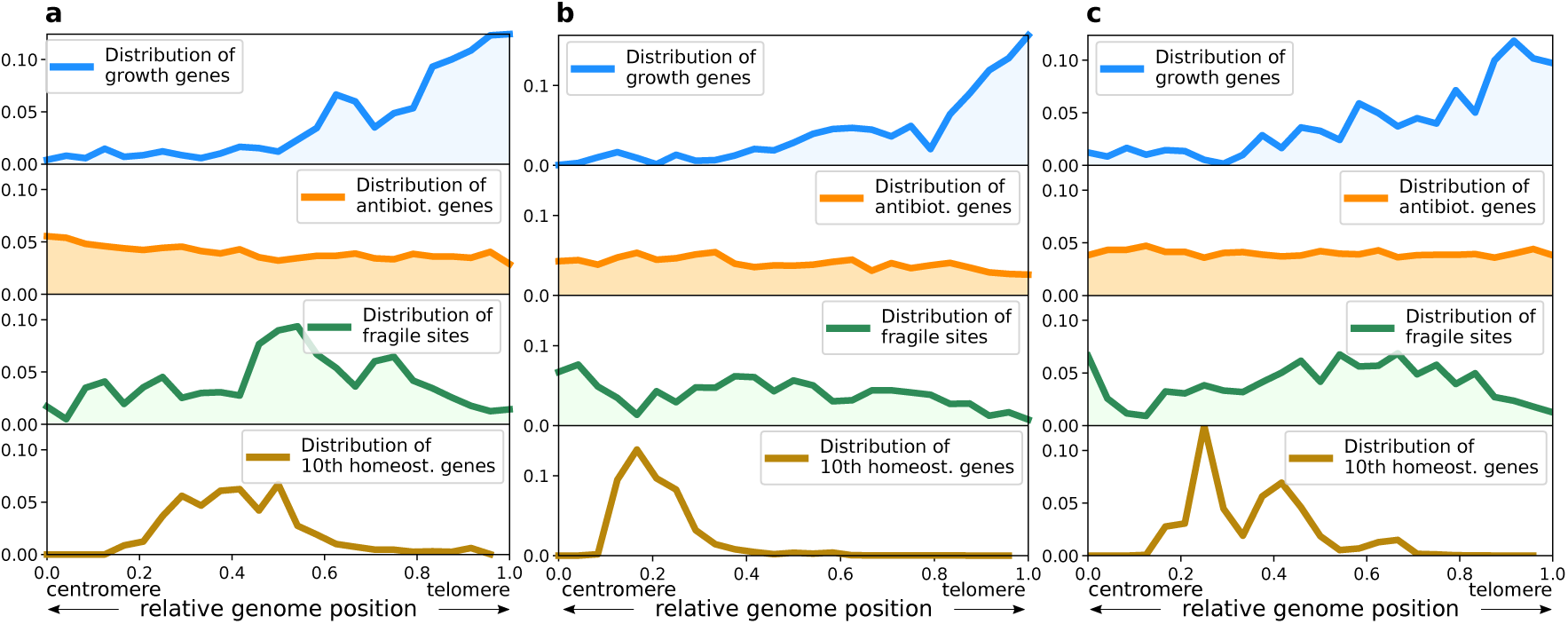
Spatial localization of genes in the genome, for three runs (**a, b, c**) in which an additional gene type is present. The first three rows in all pane show that normalized genomic location of growth-promoting genes, antibiotic genes and fragile sites. The last row shows the distribution of the genomic location of the 10th homeostatic gene - since at least *n*_*h*_ = 10 copies must be present for survival.

### S11 Division of labor evolves if colonies have sufficient time to develop and compete

At the beginning of each growth cycle, bacteria replicate locally and expand into the available space. During this initial phase, bacteria with more growth-promoting genes are at an advantage. When we reduce the duration of the growth cycle we observe that selection favors growth over antibiotic production (Fig. SF13, cycle duration *<* 1000 time steps). Genomes accumulate growth-promoting genes but no antibiotic genes - and consequently do not divide labor. When growth cycles are further reduced (*≤* 500 time steps), population size does not recover between sporulation events (which sample a fraction of bacteria) and the system goes extinct. With longer growth cycles (*≥* 1000), colony development progresses to the point that interference competition between colonies becomes common, because colonies come in contact with one another. This selects for antibiotic production (Fig. SF13), which, in turn, selects for division of labor and a genome architecture that makes this possible. In summary, inter-colony competition drives the evolution of antibiotic diversity and genome instability, and determines the condition for the emergence of division of labor.

**Figure SF13:**
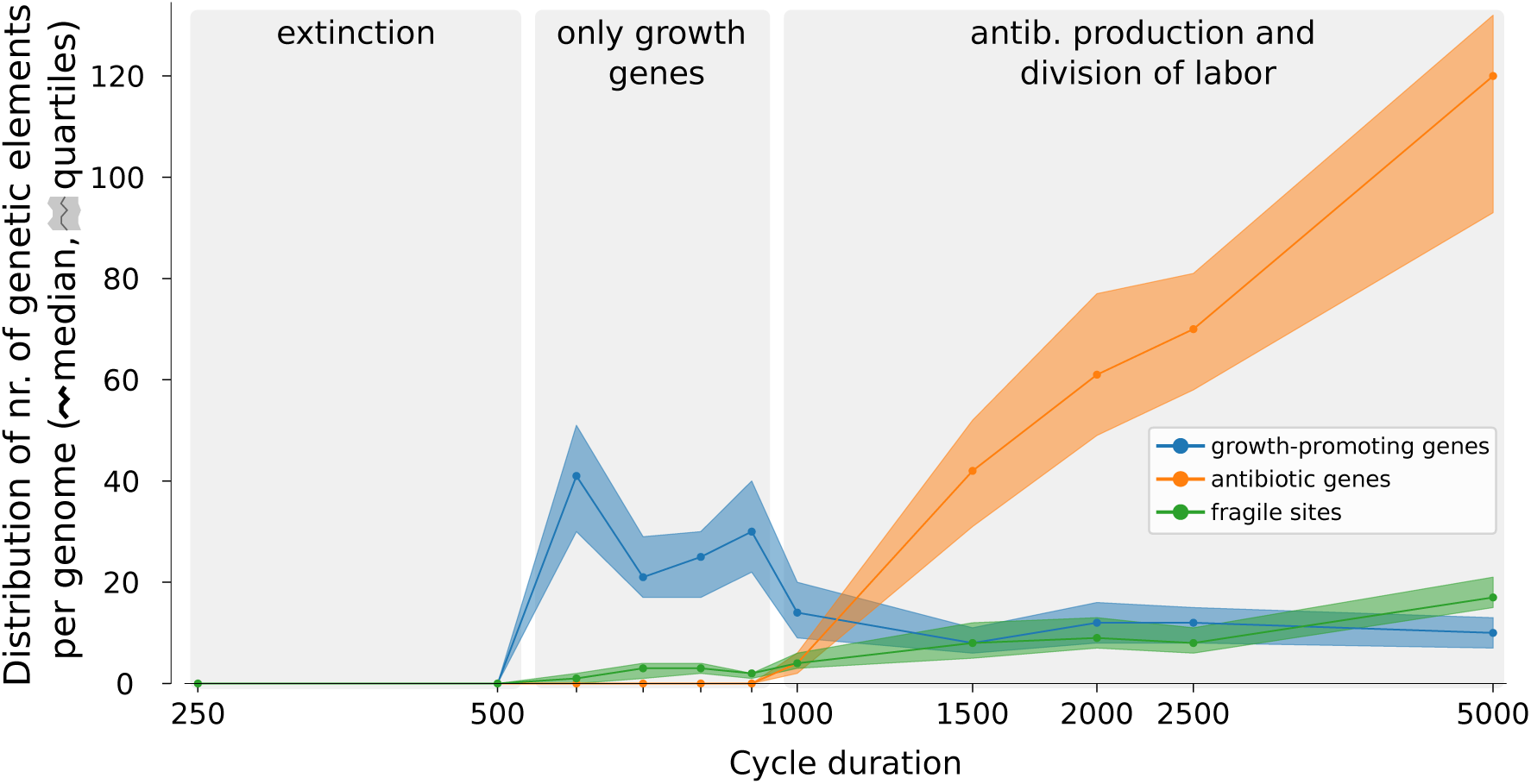
The genome composition depends on the duration of the growth cycle. The plot shows the distribution of growth-promoting genes (dot: median, shaded area: quartiles), antibiotic genes and fragile sites in the genomes of all individuals in a population, after long-term evolution, for different durations of the growth cycle (for cycle duration *≤* 500 the system goes to extinction).

### S12 The effect of destroying spatial structure

Starting from an evolved colony, we ran five simulations in which the location of bacteria was randomized at every time step. This disrupts colony formation, with two consequences: all bacteria are likely exposed to all antibiotics (in a colony this is not the case, because bacteria at its center do not come in contact with antibiotics other than those they themselves produce), and the local benefit of antibiotic production is also lost. We find that division of labor is maintained and the number of antibiotic genes increases, presumably because non-local competition always selects against loss of antibiotic resistance and favors an increase in antibiotic diversity. Fig. SF14 shows that this results in a steady increase in the number of antibiotic genes.

**Figure SF14:**
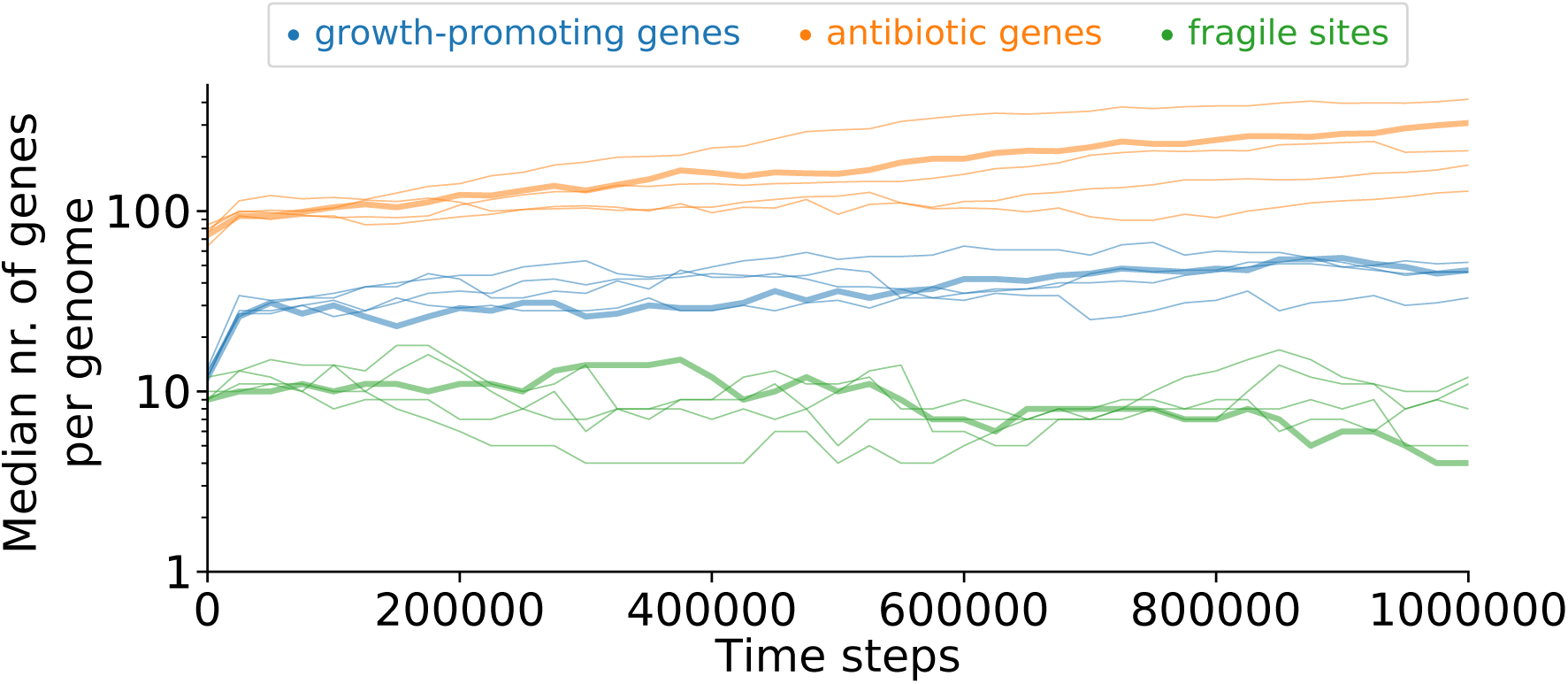
Starting from genomes that divide labor, evolutionary dynamics of the genomes in the population when the system is mixed every time step. The system is initialized from evolved bacteria that divide labor. The plot shows the median number of each gene type in the genome. The lattice location of each bacterium is randomized every time step. All the runs are initialized from the same evolved colony.

We also tested the effect of disrupting spatial structure on the evolution of bacteria that do not divide labor, i.e. starting from randomly generated genomes. Fig. SF15 shows that genomes evolve a large number of growth-promoting genes, and no antibiotic genes or fragile sites, indicating that they are solely being selected on the basis of their growth rate.

**Figure SF15:**
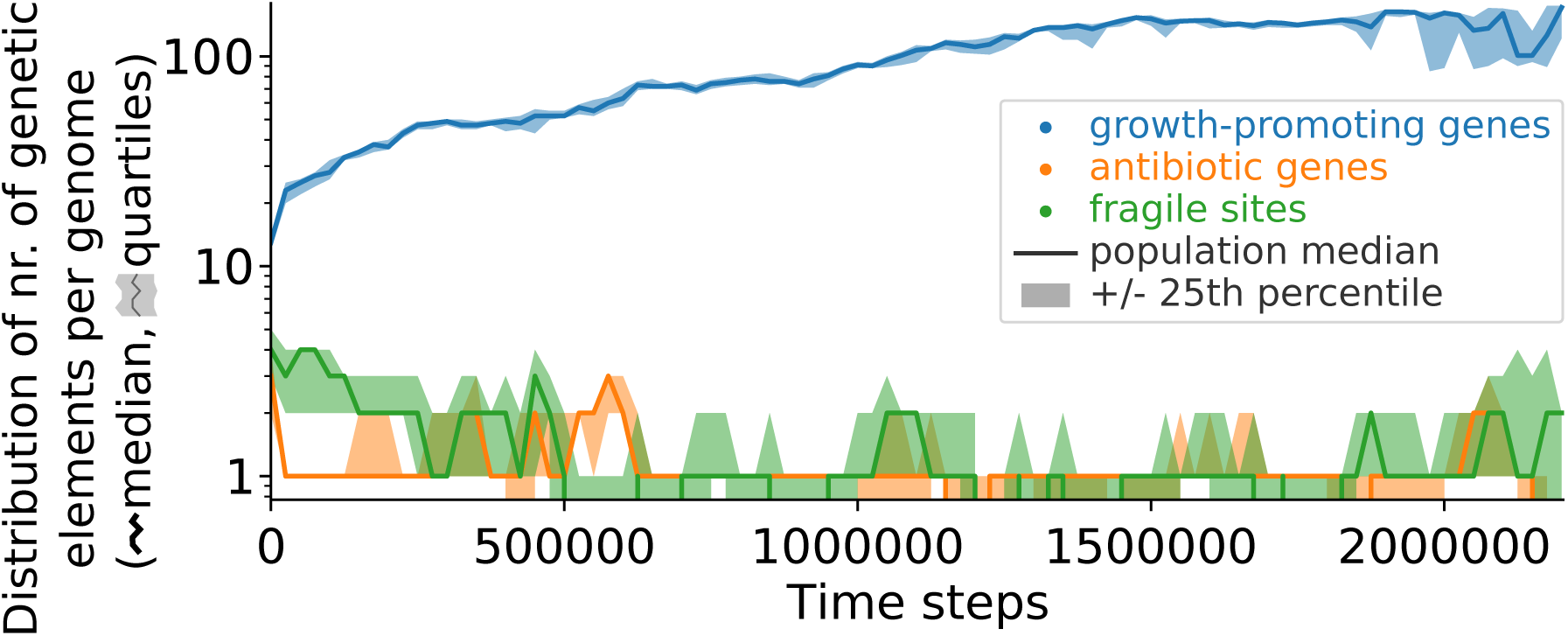
Starting from random genomes, evolutionary dynamics of the genomes in the population when the system is mixed every time step. The simulation is initialized from a population of random genomes of length 20, with average proportion of growth-promoting genes, antibiotic genes and fragile sites 12:4:4. The plot shows the median and quartile values of the number of each type of genetic element, calculated from all genomes in the population, at the end of every growth cycle.

### S13 The number of antibiotic genes is due to selection for diversity

To check that antibiotic diversity is under selection rather than the number of antibiotic genes, we modified the antibiotic production rate so that it is independent of the number of antibiotic genes. In the modified system (cf. Methods), antibiotic production per unit time 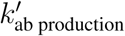 depends solely on the number of growth promoting genes - if at least one antibiotic gene is present, and is zero otherwise:

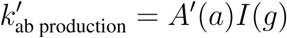

with

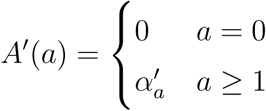

and *I*(*g*) = *exp*(−*β*_*g*_*g*) as in the Methods. Fig. SF16 shows that results are robust to this change, and a large number of antibiotic genes is incorporated in the evolved genomes, indicating that selection is on antibiotic diversity rather than antibiotic number. (cf. Fig. SF2).

**Figure SF16:**
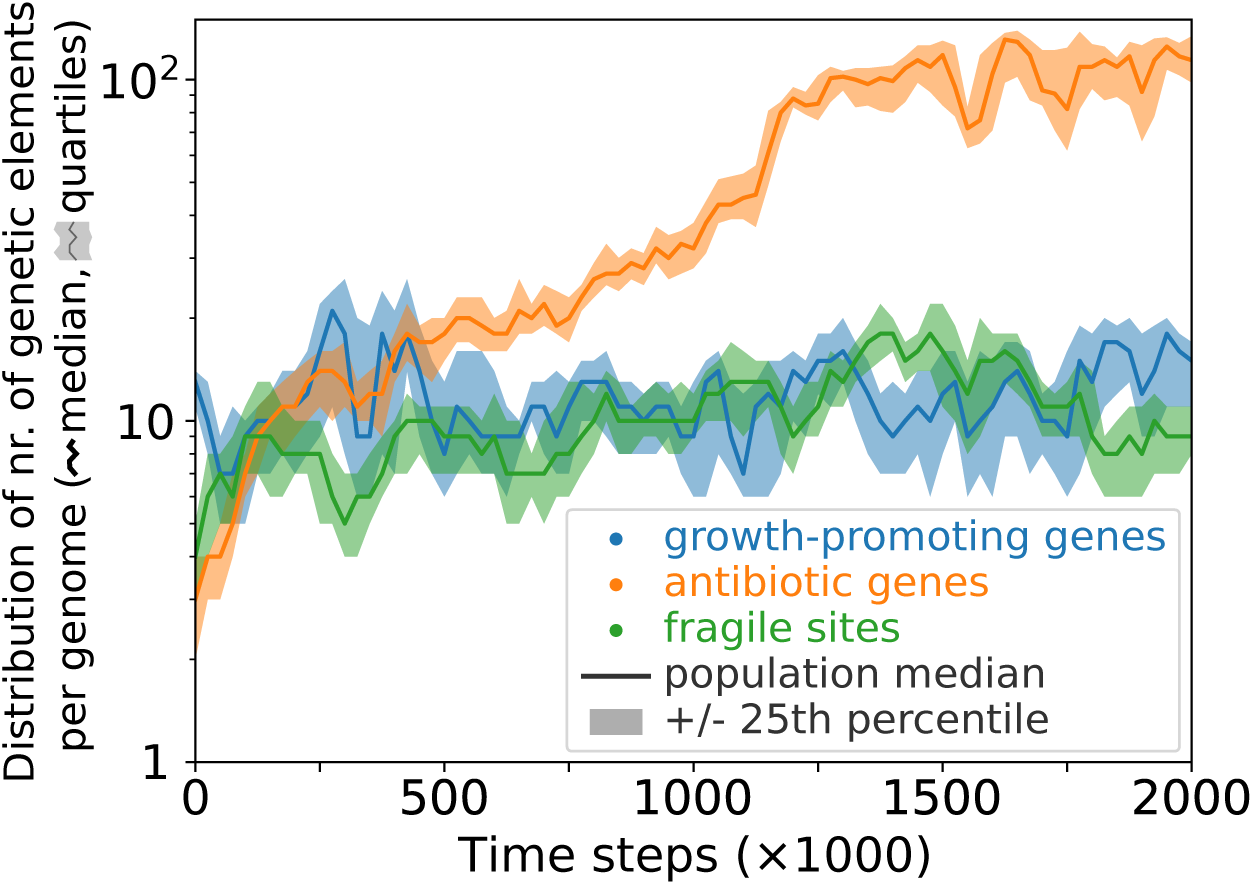
A large number of antibiotic genes is incorporated in the genome despite the antibiotic production rate being independent of the number of antibiotic genes. The plot shows the median and quartile values of the number of each type of genetic element, calculated from all genomes in the population, at the end of every growth cycle. All other parameters are identical to those of the simulation shown in Fig. SF2.

### S14 High and diverse antibiotic production

The large number of antibiotic genes and their variability are due to selection for antibiotic diversity (multi-toxicity), see Fig. SF2. Fig. SF17 shows that colonies are susceptible to most antibiotics.

**Figure SF17:**
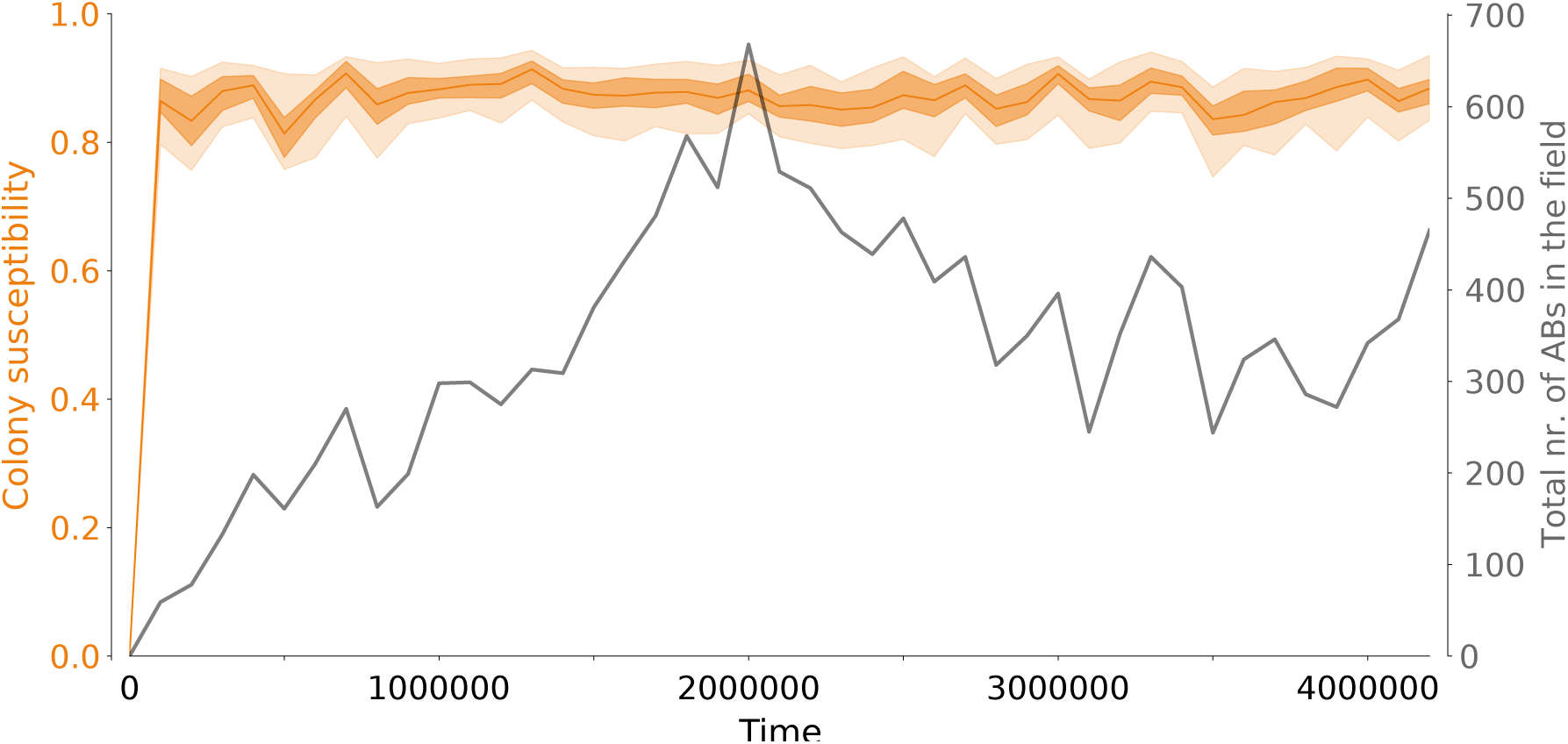
Evolutionary dynamics of antibiotic production and susceptibility. We extracted the total number of antibiotics produced per time step from the same run shown in Fig. SF2, and measured the susceptibility of each bacterium in the lattice as the fraction of antibiotics that causes a 75% fitness decrease or more. Orange line: median susceptibility, shaded areas are, top to bottom, 5th, 25th, 75th and 95th percentile. The gray line shows the total number of different antibiotics in the system.

### S15 The total number of possible antibiotics determines the evolution of colony susceptibility

The evolutionary potential for antibiotic diversification depends on the total number of possible antibiotics, which is determined by the length of the bit-string that defines the antibiotic (Fig. SF18; with a binary strings of length *ν*, the volume of the antibiotic space is 2^*ν*^). With long antibiotic strings (*ν ≥* 8), bacteria occupy a small part of the total antibiotic space, and bacteria are susceptible to most antibiotics produced by other colonies (Fig. SF18 red line, see also Suppl. Section S14). This indicates that large antibiotic diversity promotes competition - an eco-evolutionary outcome previously named “multi-toxicity” in the context of colicin evolution models [36]). When the evolutionary potential for antibiotic diversification is small (*ν ≤* 6), the system reaches a different eco-evolutionary steady state - characterized by low susceptibility because each genome contains many copies of each possible antibiotic gene. This state has been previously called “hyper-immunity” and persists because the loss of resistance to any antibiotic leads to extinction.

**Figure SF18:**
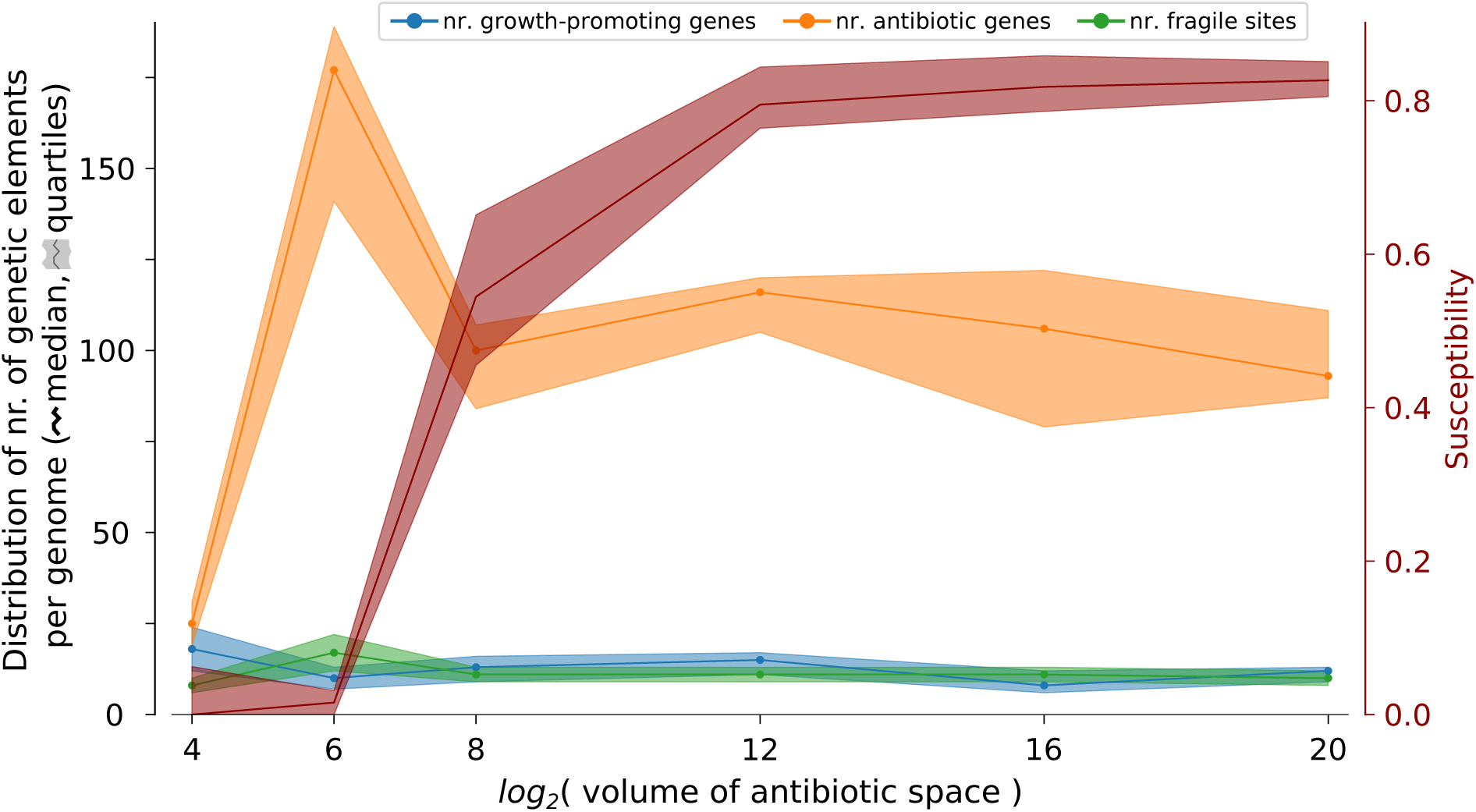
The total number of possible antibiotics, i.e. the volume of the antibiotic space, determines steady-state genome size and colony immunity. We ran one simulation for each antibiotic bit-string length *ν* ∈ 4, 6, 8, 12, 16, 20. The volume of the antibiotic space is 2^*ν*^. The plot shows the distribution of growth-promoting genes (dot: median, shaded area: quartiles), antibiotic genes, fragile sites in the genomes of all individuals in a population, as well as the distribution of colony susceptibility (defined as the fraction of all antibiotics in the lattice to which the colony is susceptible), after long-term evolution.

### S16 The architecture of genomes evolved when antibiotic volume space is small

We ran a simulation where the size of the antibiotic bit-string was *ν* = 6. After long-term evolution we measured the genome architecture in the population in the same way as shown in Suppl. Section S8. Fig. SF19 shows that the volume of the antibiotic space does not affect the evolution of the architecture that supports division of labor.

**Figure SF19:**
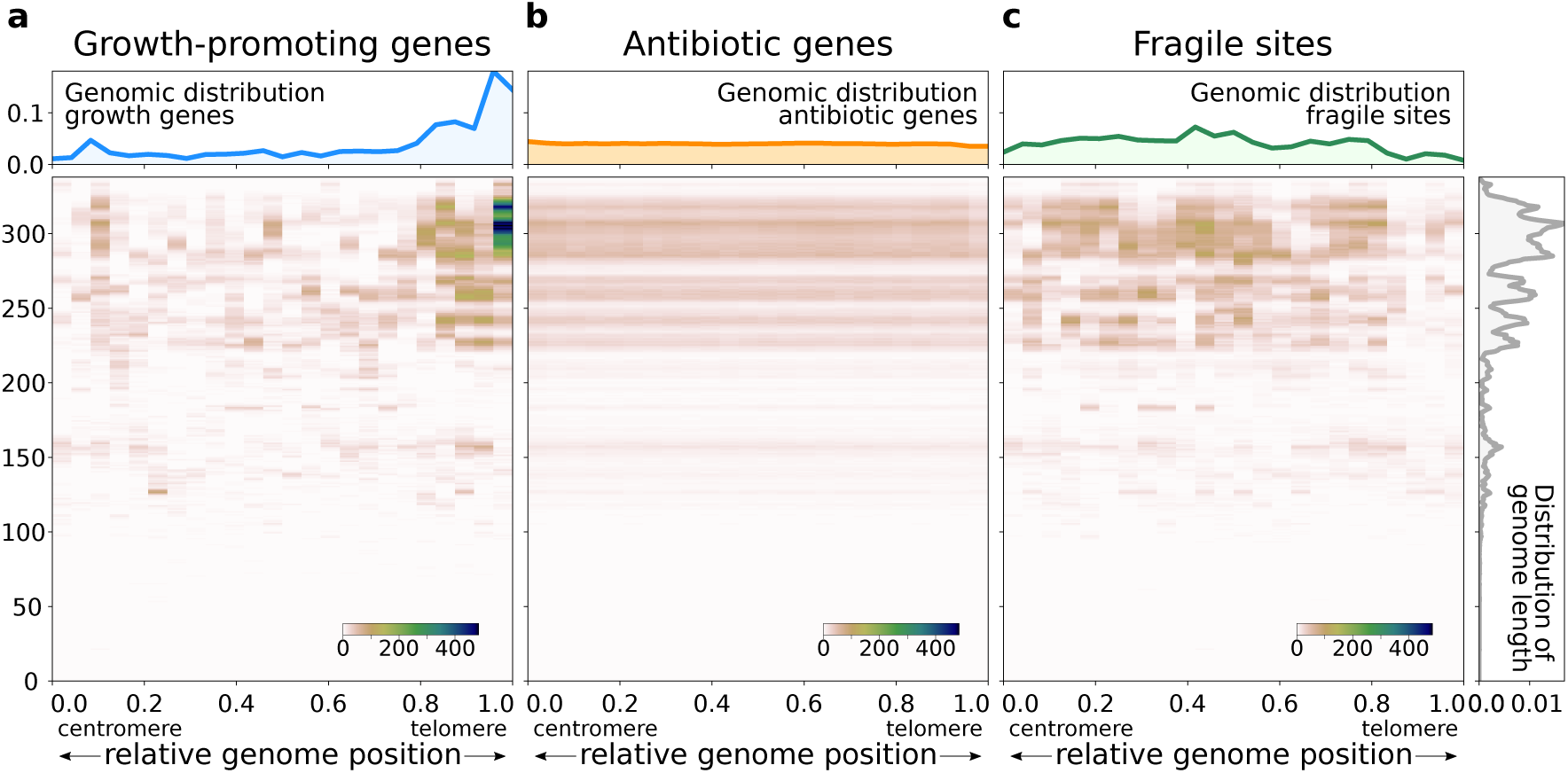
Small antibiotic space (2^6^ = 64 different antibiotics) volume does not affect division of labor. We ran a simulation identical to that shown in Fig. SF2, except for an antibiotic bit-string length of *ν* = 6.

### S17 Division of labor evolves when the deposition zone of antibiotics is smaller and when the resistance to antibiotics is broader

#### Smaller deposition zone of antibiotics

We ran a series of simulations in which we varied the radius of the circle in which antibiotics are deposited by bacteria. We ran two simulations for the following values of the radius: *r*_*α*_ ∈ *{*1, 2, 3, 5, 8*}*, and we included the results of the five simulations shown in Suppl. Suppl. Section S1 which have default radius *r*_*α*_ = 10. Fig. SF20 shows that antibiotic genes, fragile sites and growth genes are accumulated in genomes that divide labor when *r*_*α*_ *≥* 3 (although for *r*_*α*_ = 3 this happens only in one of the two simulations). Bacteria only maximize growth rate for smaller values of *r*_*α*_ antibiotics, and neither fragile sites nor antibiotic genes are accumulated in the genome.

**Figure SF20:**
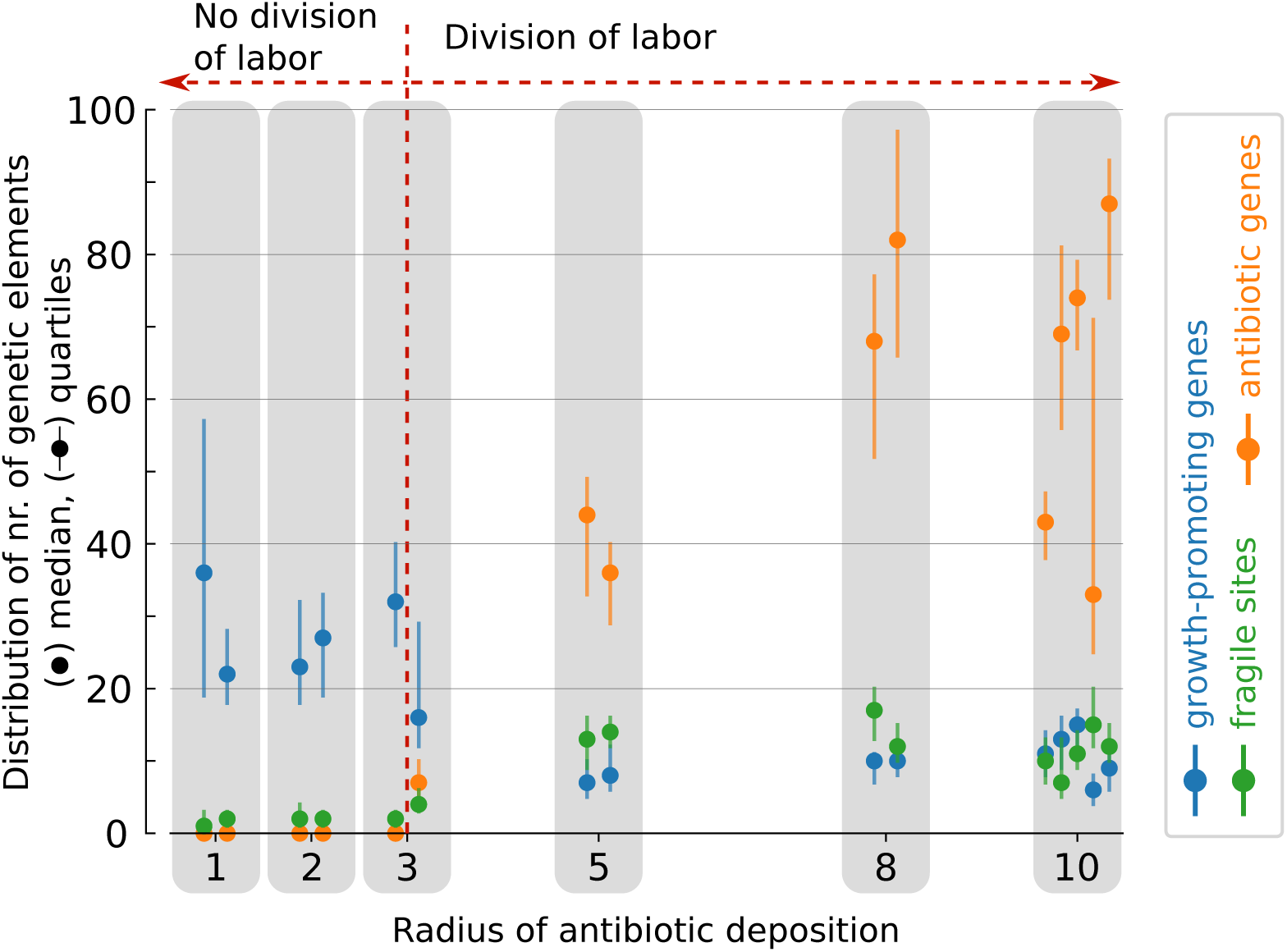
Evolutionary outcome of changing the radius of antibiotic deposition. Two simulations per radius length are run and the distribution of the nr. of genetic elements is reported, except for radius = 10, for which the five runs from Suppl. Section S1 are used. Data collected after long term evolution, at the end of a growth cycle. All other parameters and initialization are the same as in Suppl. Figure SF1.

#### Broader resistance to antibiotics

Antibiotic resistance in the model is determined by matching the antibiotic bitstring with the bitstring of the antibiotic genes. A larger difference between the two strings results in a lower resistance of the bacterium, according to the function 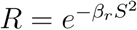 (see Methods). A higher value of *β* results in a lower resistance even for small mismatches, whereas a smaller *β*_*r*_ makes resistance more broad. We ran a series of simulations in which we varied *β*_*r*_ (Fig. SF21). Division of labor evolves when *β*_*r*_ *≥* 0.0003, indicating that the system is extremely robust to decreasing benefits of antibiotic production (for comparison, the value used for main text results is *β*_*r*_ = 0.3). For very small *β*_*r*_, any antibiotic gene provides resistance to any antibiotic, and thus there is no benefit from secreting antibiotics. The system does not evolve division of labor in this case, and instead maximizes growth.

**Figure SF21:**
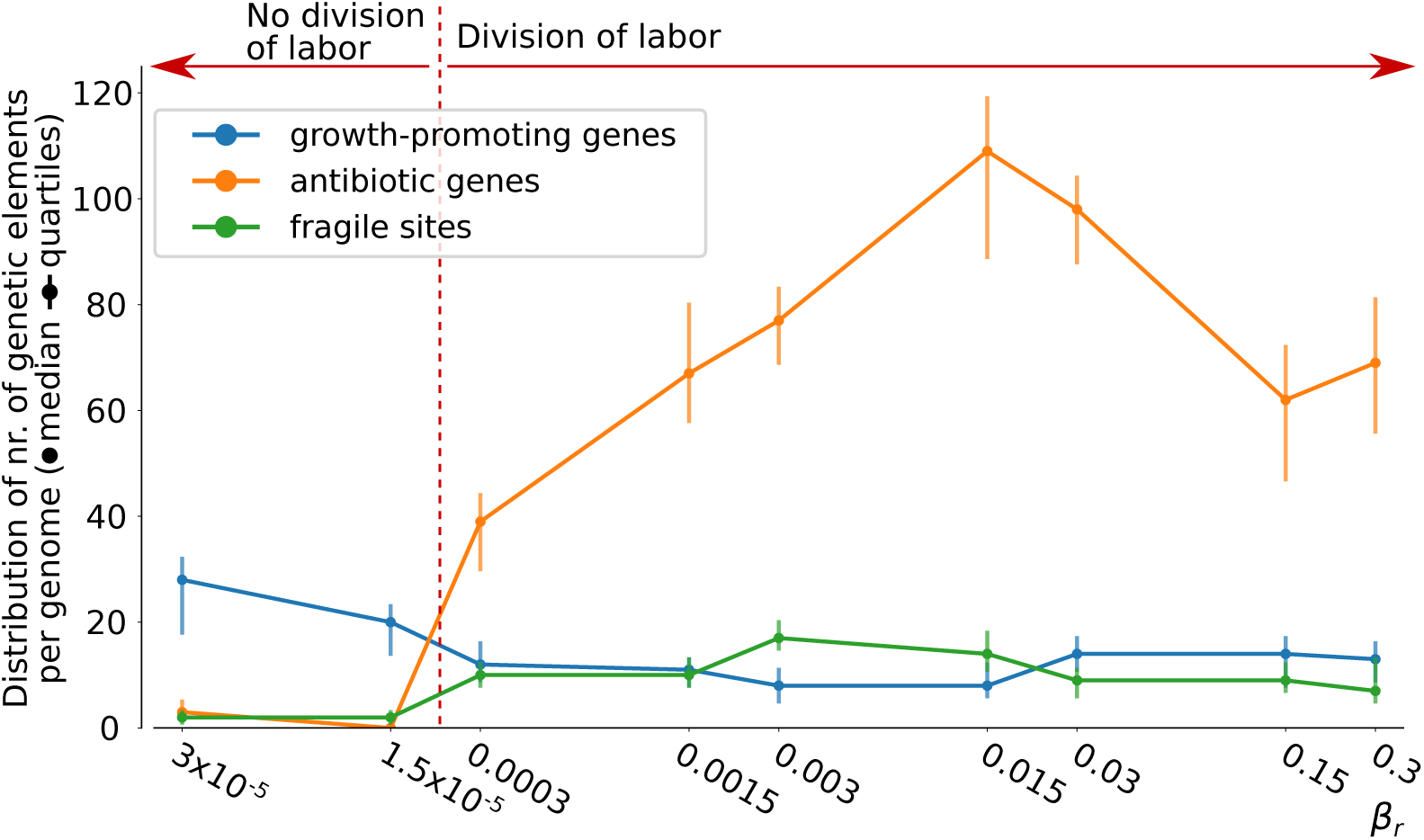
Evolutionary outcome of changing the antibiotic resistance factor *β*_*r*_. One simulation for each value is run, and the distribution of the nr. of genetic elements is reported. *β*_*r*_ = 0.3 is the value used for the simulations presented in main text. Division of labor can be inferred from the increase in number of antibiotic genes and fragile sites for *β*_*r*_ *≥* 0.0003. Data collected after long term evolution, at the end of a growth cycle. All other parameters and initialization are the same as in Suppl. Figure SF1.

### S18 Mutation-driven division of labor evolves over a wide range of (per-fragile sites) mutation rates

In main text Fig. 5 we show that division of labor evolves by showing the distance between the median nr. of growth-promoting genes in the genome of antibiotic producing bacteria and replicating ones. Fig. SF22 shows the distribution of antibiotic producing and replicating bacteria used to generate the the figure in main text.

**Figure SF22:**
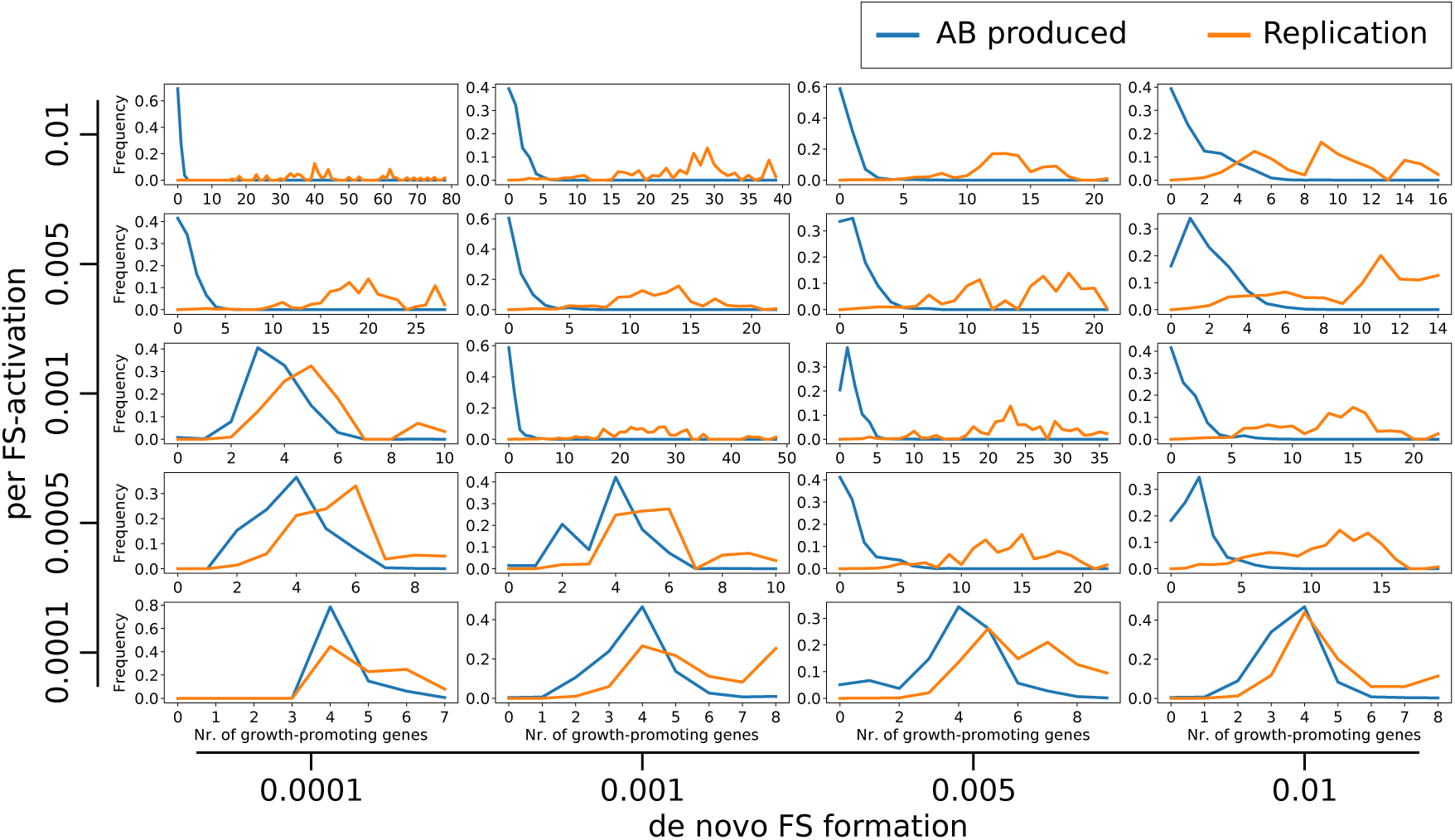
Frequency of antibiotic producers (blue) and replicating bacteria (orange) as a function of the nr. of growth-promoting genes in their genome. One simulation is run with the parameter combination indicated in the figure (all other parameters are identical to those in the caption of Fig. 1). Data is collected from the entire population, for one growth cycle after long-term evolution from two data points (in all cases after at least 600 generations). A larger difference between the two curves indicates division of labor, because the two tasks are carried by genetic distinct individuals in the same colony.

### S19 The genome composition of populations that evolve division of labor over a wide range of (per-fragile site) mutation rates

Fig. SF23 shows the genome composition in the evolved population for different value of fragile site mutation rates *μ*_*f*_ and *μ*_*n*_. A larger number of growth-promoting genes corresponds to division of labor because these genotypes cannot produce a lot of antibiotics, which are instead overproduced by other members of the colony (which arise through mutations). The same data is used to generate both this figure and Fig. SF22.

**Figure SF23:**
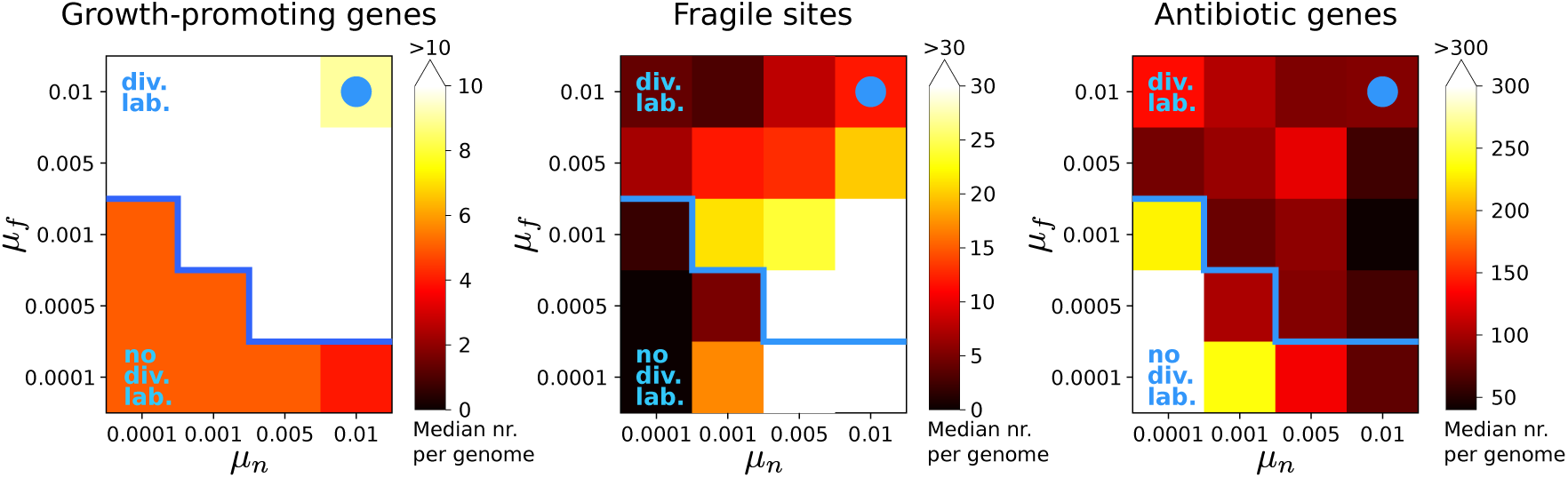
Each plot shows the median number of genetic elements - growth-promoting genes, fragile sites and antibiotic genes - per genome in a population evolved with the indicated fragile site mutation rates (the same data is used to generate Fig. SF22). The blue dot (top right corner of each heatmap) indicates default values used in the rest of the manuscript. The blue line indicates the approximate location of the boundary for the evolutionary phase transition between division of labor (above the line), and generalist (below).

### S20 Division of labor persists in evolved genomes when de-novo fragile site formation is set to zero

Division of labor persists in six independent simulations initialized with one of three evolved genomes, despite an absence of de-novo fragile site formation (i.e. *μ*_*n*_ = 0). Fig. SF24 shows genome composition after 500 *×* 10^3^ to 680 *×* 10^3^ time steps (200 to 272 growth cycles). The number of fragile sites is small but maintained above zero, and the number of both growth-promoting genes and antibiotic genes is large, indicating that division of labor occurs to enable both cell division and antibiotic production. Moreover, the evolved genome architecture also persists.

**Figure SF24:**
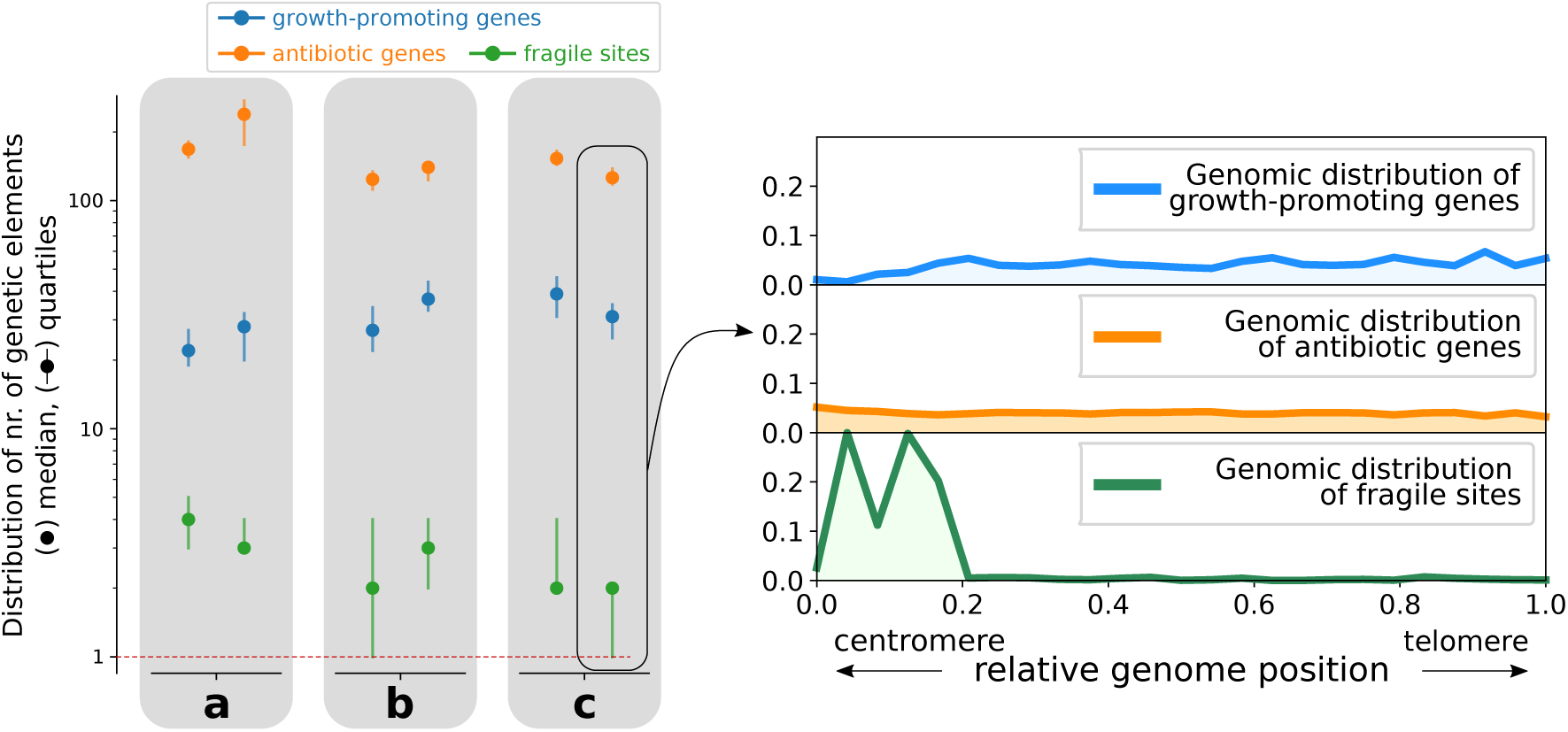
Genome composition (for all runs) and genome architecture (for one case) after long term evolution with *μ*_*n*_ = 0. The plot on the left shows the final genome composition for six simulations started from three independently evolved genomes (a,b,c). On the right, the genome architecture extracted from the whole population at the last time point of the simulation is plotted to show that resulting genomes are organized so that fragile sites are at the 5’ of growth genes, and thus can generate antibiotic-producing mutants.

